# Influenza A viruses induce tunnelling nanotube-like structures through the onset of apoptosis

**DOI:** 10.1101/2024.09.25.614890

**Authors:** Daniel Weir, Calum Bentley-Abbot, Jack McCowan, Colin Loney, Edward Roberts, Edward Hutchinson

## Abstract

As well as spreading through virions, influenza A viruses (IAVs) can evade antiviral drugs and neutralising antibodies by spreading directly from cell to cell. In cell culture this can occur by the induction of intercellular membrane connections known as tunnelling nanotube-like structures (TLSs), which are capable of trafficking the viral genome between cells. Here, we showed that TLSs are present at the site of IAV infections *in vivo*, and then used *in vitro* models to ask how IAVs induce their formation. We found that TLS induction cannot be induced by cytokine signalling from infected to uninfected cells, but requires IAV replication within cells. IAV replication can form filamentous virions with structural similarities to TLSs, but we found that TLS induction is independent of virion morphology. We therefore looked at the intracellular responses to infection. Using a pan-caspase inhibitor, we found that TLS induction by IAVs requires the onset of apoptosis. Our results, which suggest that IAVs control their ability to spread directly from cell to cell by driving infected cells into apoptosis, identifies a new way in which a virus can manipulate its host to evade antiviral immune responses.

**Author Summary:** Influenza A viruses (IAVs) spread efficiently through the respiratory tract in the form of extracellular virus particles, but can be restricted by neutralising antibodies and antiviral drugs. IAVs can avoid this restriction by transporting viral genomes directly from one cell to the next. They can do this by inducing the formation of long, thin intercellular connections known as tunnelling nanotube-like structures, which are capable of trafficking viral genomes. In this study, we demonstrate for the first time that tunnelling-nanotube like structures form within IAV infected lungs. We then asked how IAVs induce these structures. We found that cell death pathways triggered by IAV replication induce the formation of tunnelling nanotube-like structures, thereby establishing routes of infection spread to other cells. In this way, the virus exploits the cell death response of its host to ensure that its infection can continue to spread even within the restrictive environment of the respiratory tract.

## Introduction

Influenza A viruses (IAVs) require virions for transmission between host organisms. However, within an infected host IAV infection can spread between cells through alternative routes that are independent of extracellular virions (1–5). This process, referred to as direct cell to cell spread, can ensure the continued intercellular transmission of IAV infection in the presence of neutralising antibodies and antiviral drugs (1–4, 6).

Many viruses can undergo direct cell to cell spread and there are diverse mechanisms by which they do so (reviewed in (6)). Some of these transport complete virions directly from one cell to another without releasing them into the extracellular environment. For example, filopodial bridges and virological synapses can deliver newly formed virions of alphaviruses and human immunodeficiency virus (HIV), respectively, to adjacent cells through cellular junctions or synaptic clefts (7–9). Other mechanisms use membrane fusion to enable the direct transfer of viral genomes and proteins. This can be seen in the formation of syncytia from directly adjacent cells (10), as well as in connections made between distant cells through open ended, long-range connections known as tunnelling nanotubes (TNTs) (11). TNTs are thin (typically 50 to 200 nm in diameter), F-actin rich cell connections that function in trafficking various cargos (12, 13) and have been implicated in the spread of several viral pathogens (11) including herpes simplex virus (HSV), HIV, SARS-CoV-2 and IAVs (1, 14–17).

Studies of *in vitro* models have shown that IAV infection induces the formation of TNT-like structures (TLSs), creating an F-actin and Rab11a dependent mechanism for infection of cells and for the reassortment of viral genome segments (1, 3, 5). However, TNTs are delicate and transient structures (12), making their detection within tissues such as respiratory epithelia extremely challenging. Therefore, it was unclear if TLSs could physically form in the dense tissues which IAV naturally infects. It was also unclear how IAV infection could induce TLS formation.

In this study, we addressed these questions. Using a reporter mouse system, we observed TLSs forming from IAV infected cells within the lung epithelium, supporting the relevance of TLS trafficking during natural IAV infections. Using *in vitro* assays, we then asked how IAV infection induces TLS formation. Having ruled out cytokine signalling from infected to uninfected cells, as well as processes linked to the formation of virions, we found that the induction of TLSs by IAVs results from the triggering of apoptosis. The onset of apoptosis is known to drive TLS formation between healthy and apoptotic cells. By using a pan-caspase inhibitor to prevent apoptosis, we completely prevented the ability of IAV infected cells to induce the formation of TLSs. Our data therefore reveal that TLS connections between cells are a feature of natural IAV infection that are induced when viral replication triggers the onset of apoptosis.

## Results

### TNT-like structures are present within the lungs of IAV infected mice

To date, the study of TLSs and their consequences for IAV infection has been limited to *in vitro* models. However, for TLSs to be relevant to the intercellular spread of IAV infections they need to be present at the site of natural infection. Within the lung, TLSs have been observed within solid tumours (18) but not within typical lung epithelium. Conceptually, the lung epithelium is a challenging environment for cells to form long-range, thin intercellular projections. These already fragile and transient structures would need to be able to form in a dense and complex environment whilst withstanding the mechanical stresses of breathing and mucociliary flow. Furthermore, the visualisation of TLSs in tissue, either *in vivo* or *ex vivo,* has been mostly limited to tissues with lower cell densities and/or high rigidities (14, 19–21), with only some non-infection based stress or pathologies studied (e.g. lipopolysaccharide treatment (19), or within solid tumours (21–23)). Such tissues have been imaged with low complexity staining and tissue manipulation (18, 22, 23), and the study of TLS formation between epithelial cells following respiratory virus infection is challenged by the need to distinguish between infected and healthy cells.

To assess the ability of TLSs to form at the site of a respiratory infection, we needed a method which uniquely labelled infected cell membranes with minimal tissue manipulation to maintain TLS integrity prior to imaging. We did this using the mT/mG reporter mouse system in combination with an IAV (A/Puerto Rico/8/1934 H1N1) which has been genetically modified to encode the Cre recombinase (PR8 Cre) (24). The mice encode a membrane-targeted tdTomato (mT) fluorophore flanked by *loxP* sites, such that when Cre recombinase is expressed, the *tdTomato* gene is replaced by a downstream membrane-targeted *GFP* gene (mG, Fig 1A) (24). The result is that infection with a Cre-expressing virus permanently changes the membrane fluorescence of a cell. Accordingly, we infected mT/mG mice with PR8-Cre and, six days post infection, we harvested the lungs and identified mG positive infected cells using thick tissue section, super resolution confocal imaging (Fig 1B and C).

**Fig 1.**
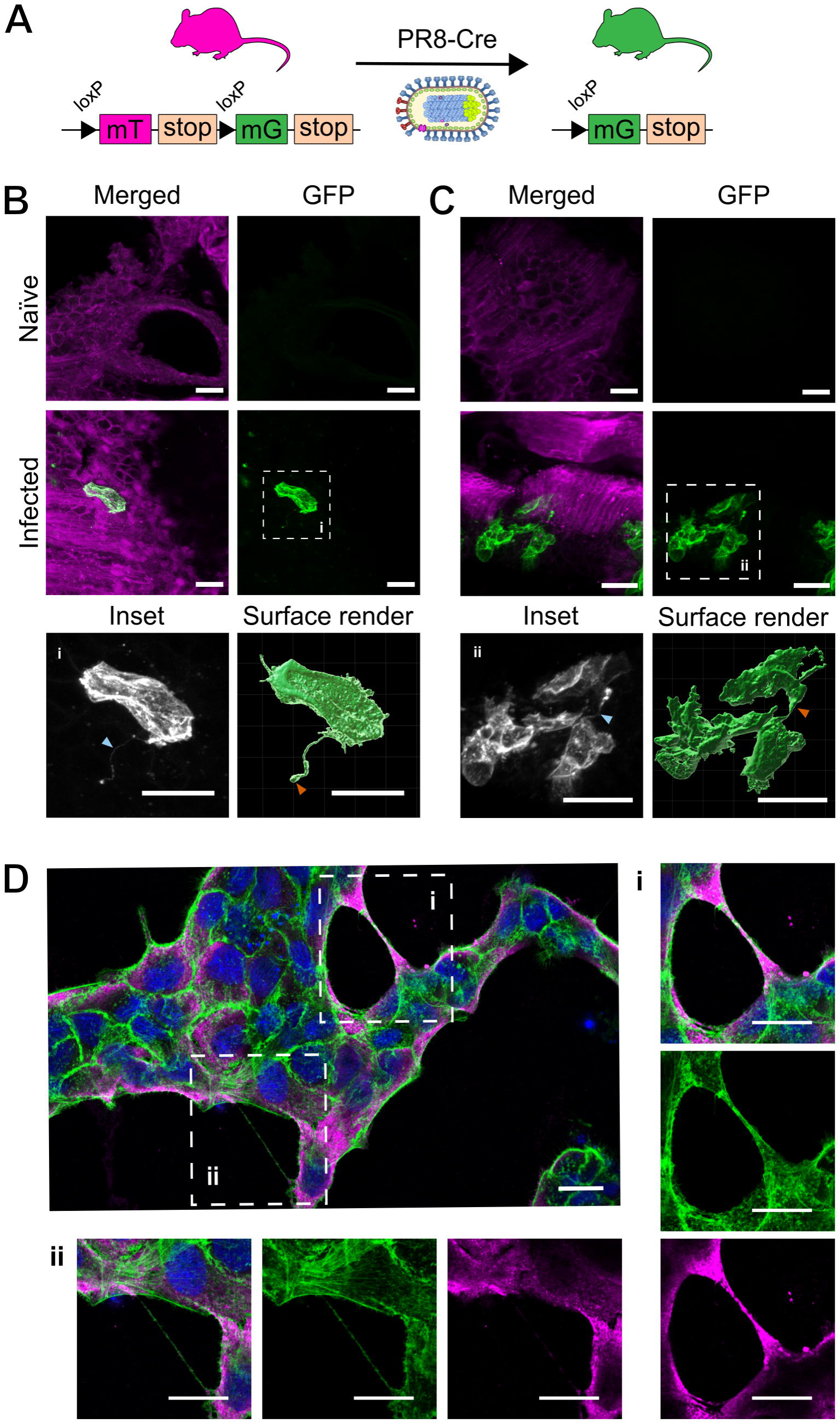
TNT-like structures involving IAV infected cells are observed *in vivo* and *in vitro*. **(A)** Schematic of mT/mG reporter mouse system in combination with an IAV encoding for Cre (PR8-Cre), with arrows indicating *loxP* sites that flank the tdTomato fluorophore. Following intranasal infection and Cre recombinase expression, the membrane targeted tdTomto (mT) is replaced with a membrane targeted GFP (mG) fluorophore. Six days after intranasal infection with PR8-Cre, lungs were harvested from mT/mG mice and thick sections were prepared for confocal microscopy. **(B & C)** Maximum intensity projections of TNT-like structures (TLSs) within lung tissue **(B)** projecting from an isolated PR8-Cre infected cell (distinguished by the expression of GFP), and **(C)** connecting infected cells. Magnified insets outlined in white boxes are shown (i-ii) alongside surface renders. Membrane labels are tdTomato (magenta) and GFP (green/white). Blue arrowheads indicate the presence of TLSs, and orange arrowheads indicate the bulbous termini of TLSs. **(D)** Tiled image of a single field of view (63x magnification), representative of images used for TLS quantification, showing TLSs of varying thickness between MDCK cells at 16 h post infection (h.p.i.) with PR8 at an MOI of 1.5 PFU/cell. Nuclei (blue), F-actin (green), nucleoprotein (magenta). All scale bars = 20 μm.

Within infected lungs, we clearly observed IAV infected cells, and from the surface of these we could observe the protrusion of very thin TLSs (Fig 1B). Interestingly, these appeared to have formed in the spaces between cells of the epithelium rather than through the lumen of the airway, resulting in a curved structure similar to the contours of adjacent cells. In addition, TLSs seemingly connecting infected cells were seen (Fig 1C and S1 Fig), suggesting that TLSs might be capable of transferring IAV infection within a tissue. The structures we observed in the airway epithelium have similar characteristics to those previously seen in mammalian tissues such as the cornea of mice, with TLSs often being curved and having a bulbous structure at the TLS terminus (19). We noted that the structures we observed were shorter than those seen in some other mammalian tissues, such as solid lung tumours (18). This would be compatible with the hypothesis that TLS length is limited in tissues with higher cell densities (19). To the best of our knowledge, this is the first demonstration of TLSs within infected but otherwise normally structured lung tissue, and the first demonstration of these structures being able to form within a layer of epithelial tissue. The presence of TLSs within tissues that undergo repetitive expansion and contraction suggests a surprising degree of robustness compared to the properties of these structures *in vitro* (12).

Having demonstrated that TLSs do form at the natural site of infection, we required an easily manipulable *in vitro* system in which we could study the interplay between IAV infection and the triggering of TLSs. By applying an optimised fixation method (25) to MDCK cells infected by IAV we were able to preserve TLSs of various thicknesses and presumed stabilities (Fig 1D). Consistent with previous cell culture based studies, we found that infected cells typically produce straight TLSs that contain punctate viral nucleoprotein (NP), indicating the incorporation and transport of the viral ribonucleoprotein complex (Fig 1D) (1, 3, 5). This demonstrated the suitability of our *in vitro* approach for the study of TLS formation following IAV infection.

### TNT-like structures are formed by both infected and uninfected cells, with no evidence of pathfinding

Whilst the mT/mG reporter mouse system revealed that TLSs were able to extend from infected epithelial cells *in vivo*, the tendency of these structures to originate or be received by infected and uninfected cells was unknown. Such a preference would suggest the occurrence of TLS pathfinding, which describes a process by which TLSs are guided from a TLS donor cell, along a concentration gradient of a secreted factor produced by the TLS recipient cell (26). Pathfinding has been demonstrated to occur between homotypic astrocyte cultures and cocultures with HEK293 cells, along a concentration gradient of the extracellular small protein S100A4 (27). In this way TLS pathfinding determines which cells form TLS connections and have functional consequences. For example, diseased cells lacking functional lysosomes induce TLSs to facilitate the transfer of lysosomes to them from healthy cells (28, 29).

In the context of virus infection, evidence of TLS pathfinding varies and appears to be virus specific. For example, TLSs have been reported to originate at comparable rates from HTLV-1 positive and negative cells in a co-culture (30). In other cases directional TLS formation has been observed, for example during vaccinia virus infection – though in this case it was driven by vaccinia virus proteins on cell surfaces which upon interacting with extracellular secondary incoming virus lead to the triggering of TLSs towards uninfected cells (31). Here, we asked whether TLS pathfinding would occur during IAV infections, as this would promote the cell to cell spread of IAV.

IAV infection results in the production of a wide variety of extracellular signalling molecules, such as interferon (IFN; reviewed in (32, 33)), that can trigger an antiviral response in neighbouring cells. We hypothesised that a potential TLS chemoattractant could be amongst the paracrine innate immune signals produced by IAV infected cells. If this was the case, we would expect to observe TLSs preferentially forming from uninfected to infected cells. To investigate this, we modified A549 cells to constitutively express a membrane targeted AcGFP fluorophore. These cells retained the membrane label upon passaging and when cocultured with WT A549s this allowed the membrane of origin of TLSs to be clearly identified. We infected the co-culture at a low MOI and determined the recipient and donor cells of the TLSs that formed (Fig 2A). We first observed that modified A549s could be infected by IAV and form TLSs at rates comparable to the WT A549s (S2 Fig). We then calculated the proportion of uninfected and infected cells producing a TLS, and found no significant difference between them (Fig 2B). Infection status also did not influence the mean TLS length (approximately 9700 nm; Fig 2C). Therefore, infection status of A549 cells did not determine their ability to initiate a TLS, or the length over which interactions could form.

**Fig 2.**
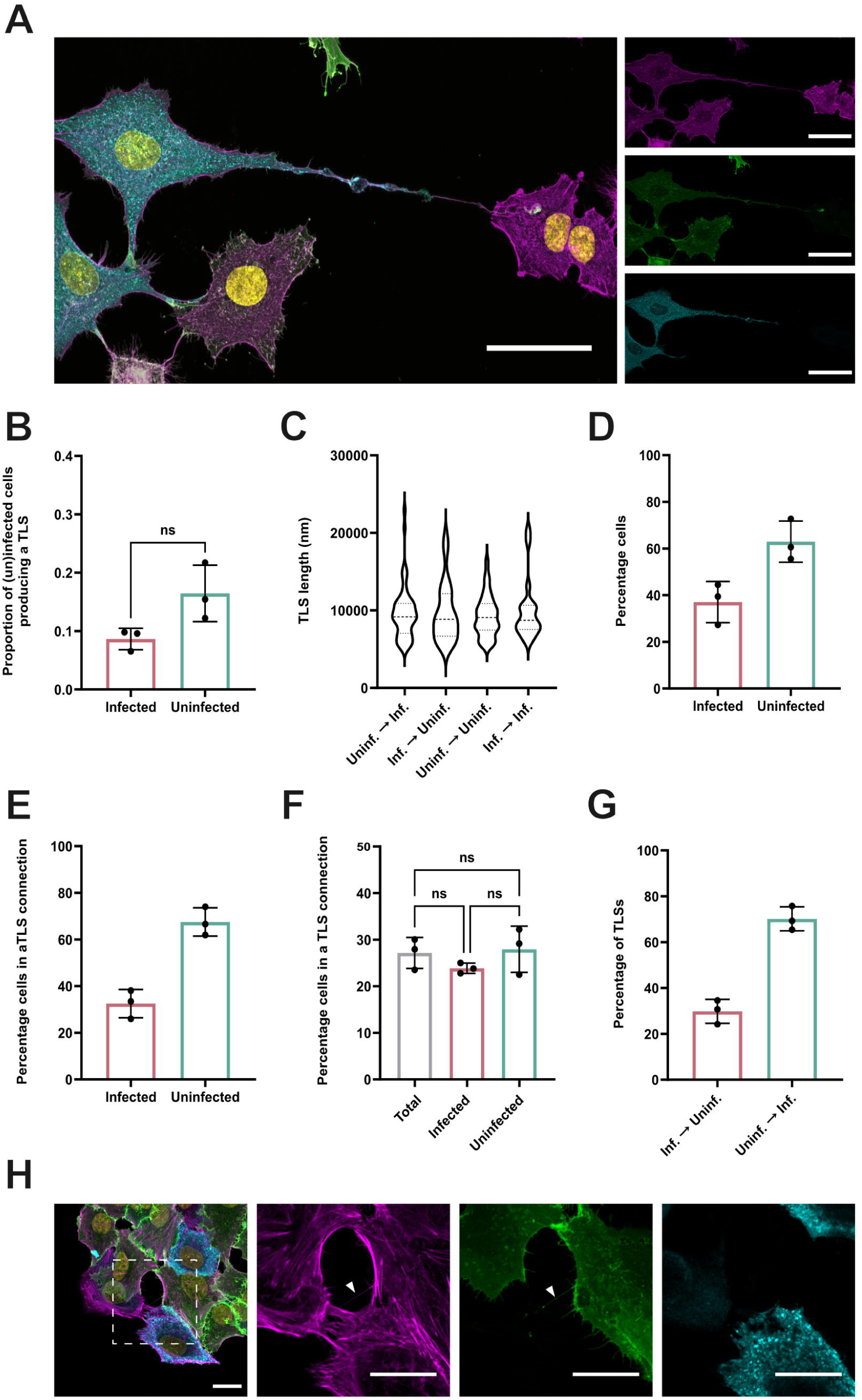
Infected and uninfected A549 cells both initiate and are contacted by TLSs at similar frequencies in co-cultures. **(A)** A TLS structure forming from a PR8 infected, AcGFP membrane labelled A549 cell, connecting to an uninfected, WT A549 cell. Nuclei (yellow), F-actin (magenta), AcGFP (green), NP (blue). Scale bar = 50 μm. **(B)** The proportion of uninfected and infected cells producing a TLS in a low MOI infection. The difference between the proportions of uninfected and infected cells that produced a TLS was tested for significance by Mann-Whitney test (n.s. *p* > 0.05). **(C)** The length of TLSs connecting cells with either an asymmetric or symmetric infection status, shown as violin plots with the median and the upper and lower quartile values indicated by dashed lines. Differences in mean TLS length was tested for significance by Kruskal-Wallis test (n.s. *p* > 0.05). **(D)** The percentage of co-cultured cells that were infected or uninfected and **(E)** the percentage of cells of infected or uninfected cells involved in a TLS connection. **(F)** The percentage of total cells, total infected cells and total uninfected cells involved in a TLS connection. Differences between infection status were tested for significance by Kruskal-Wallis test (n.s. *p* > 0.05)**. (G)** Percentage of TLSs originating from an infected or uninfected cell, collected from connected pairs of WT and AcGFP membrane labelled A549 cells (i.e. heterotypic cell pairs) with an asymmetric infection status. For all data, the mean and SD is shown (n = 3). **(H)** Representative confocal image of a heterotypic cell pair displaying asymmetric infection status, connected by a TLS originating from the uninfected cell. Nuclei (yellow), F-actin (magenta), AcGFP (green), NP (blue). Scale bar = 20 μm.

These observations suggested that the likelihood of uninfected and infected cells being involved in a TLS connection would be proportional to their abundance. According to NP staining, 37 % of cells in the co-culture were infected and 63% uninfected (Fig 2D). This was reflected in the proportion of cells involved in a TLS that were infected (Fig 2E), which suggests that TLSs were not preferentially originating from or contacting infected or uninfected cells. This is further supported by the observation that a similar percentage of total uninfected and infected cells were involved in a TLS (Fig 2F). We then focused on TLSs connections between heterotypic cell pairs (WT and AcGFP labelled) which displayed an asymmetric infection status (i.e. infected and uninfected), as this allowed us to correlate TLS directionality to the infection status of donor and recipient cells. We observed that approximately 70 % of TLSs mediating these cell connections originated from the uninfected cell (Fig 2G & H), which once again was consistent with the proportion of uninfected cells in the assay (Fig 2D). Together, these data indicate that, under these *in vitro* conditions, IAV infection does not lead to TLS pathfinding.

### IAVs induce TNT-like structures through viral replication but not by the production of secreted factors

The absence of TLS pathfinding suggested that the secreted factors produced in response to IAV infection are not responsible for the induction of TLSs by IAV which has previously been observed (1, 3, 5). However, outside the context of infection, IFN-α treatment has been shown to drive TLS induction within Kcl-22 cells (34). Additionally, conditioned media from macrophages has been shown to induce TNTs within the breast cancer cell line MCF-7, potentially by the secretion of paracrine cytokines and chemokines (35, 36). Therefore, whether paracrine innate immune signals had a role in TLS induction in the context of an infection was unknown and required investigation.

To examine this, we used MDCK cells due to their high permissiveness to IAV infection, their ability to produce innate immune signals, and their previous use for the study of TLS induction (1, 37). We collected the medium from MDCK cells 16 h after infection with the influenza strain Udorn (at which point abundant viral replication has occurred and TLSs can be observed, Fig 1D) and inactivated virions by U.V. treatment before overlaying onto fresh cells (Fig 3A). Sixteen hours later, we used immunofluorescence to confirm virus inactivation and to quantify the number of TLSs that had formed (Fig 3Ai). Cells overlaid with U.V. treated supernatant lacked any viral NP expression, confirming that virus inactivation had been effective (Fig 3B). This correlated with a lack of TLS induction, with TLS formation being comparable to that in cells treated with media from mock infections (Fig 3C). TLS induction only occurred when additional, replication competent virus was added to the cells (Fig 3B iii and C). To test whether U.V. treatment could have inactivated innate immune signals in the conditioned medium, we harvested cell lysates and used immunoblotting to detect phosphorylated STAT1 (pSTAT1), a signalling molecule phosphorylated during type 1 IFN signalling (Fig 3A ii). We observed that when U.V. treated conditioned media taken from an infection was added to cells this doubled pSTAT1 abundance (Fig 3D & E). This response demonstrates that U.V. treatment did not prevent the conditioned medium from carrying innate immune signalling molecules, and therefore shows that these molecules were not responsible for TLS induction.

**Fig 3.**
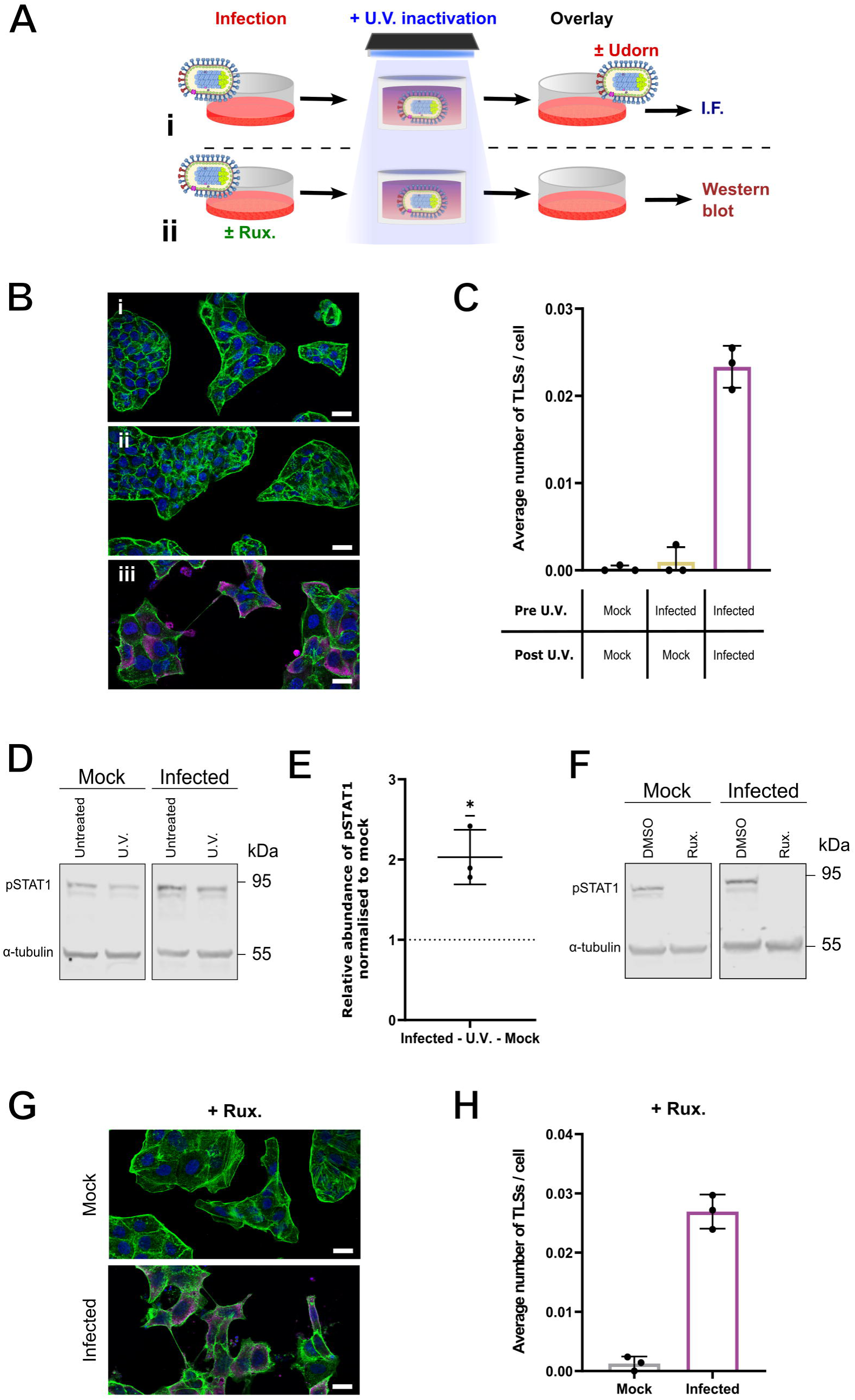
Replicating IAVs, not paracrine signals, are required for TLS induction. **(A)** Schematic of experimental design, showing the U.V. treatment of conditioned media collected from either mock or Udorn infected cells at 16 h.p.i., overlaid onto fresh cells for either downstream **(i)** immunofluorescent (I.F.) staining and confocal imaging or **(ii)** cell lysate harvesting and western blotting. **(B)** Representative confocal images of cells treated with the following overlays: **(i)** U.V. treated supernatant from mock infected cells, **(ii)** U.V. treated supernatant from infected cells and **(iii)** U.V. treated supernatant from infected cells, and additional Udorn virus. Nuclei, (blue), F-actin (green), NP (magenta). Scale bar = 20 μm. **(C)** Average number of TLSs per cell for MDCK cells at 16 h following treatment with the overlay media described. **(D)** Western Blots for pSTAT1 using cell lysates harvested 16 h post treatment with U.V. inactivated conditioned media from either mock infection or Udorn infection. Images are representative of three biological repeats. **(E)** Ratio of the relative abundances of pSTAT1 at 16 h post treatment with U.V. inactivated supernatants from infection, normalised to mock infection. No change is indicated by a dashed line; the significance of the difference from this was tested by a one-sample t-test (\**p* < 0.05). **(F)** Western blot of pSTAT1 from cells infected with Udorn at an MOI of 1.5 PFU/cell with the addition of either DMSO or 2 µM ruxolitinib (Rux.) to the SFM overlay, with cell lysates being harvested 16 h.p.i. **(G)** Representative confocal images of MDCK cells at 16 h.p.i. with Udorn at an MOI of 1.5 PFU/cell in the presence of 2 µM Rux. Nuclei (blue), F-actin (green), NP (magenta). Scale bar = 20 µm. **(H)** Average number of TLSs per cell in the presence of Rux. following mock or Udorn infection. For all data the mean and SD are shown (n = 3).

To further examine the hypothesis that TLS induction was not caused by innate immune signalling, we treated MDCK cells with 2 µM ruxolitinib, a broad-spectrum Janus kinase (JAK) inhibitor which prevents the phosphorylation of STAT1 (Fig 3F). Despite the inhibition of interferon signalling, IAV infection still clearly induced the formation of TLSs (Fig 3G & H), at levels comparable to those previously seen (Fig 3C). Together these results strongly suggest that TLS induction during IAV infection is independent of secreted extracellular factors, including the production of cytokines that signal through the JAK/STAT1 pathway.

### IAVs induce TNT-like structures and undergo direct cell to cell spread independent of virion morphology

As the induction of TLSs was dependent on viral replication, we reasoned that it might be caused by some feature of the viral replication cycle, and we looked to see if differences in the level of TLS induction between different strains of IAV could help to understand this. In particular, we noted a striking similarity between the filamentous virions formed by some IAVs and the physical features of TLSs.

IAV virions are pleiomorphic, with morphologies ranging from spherical to filamentous (38, 39). Although the most widely-used laboratory strains of IAVs have lost their ability to form filamentous virions, clinical and veterinary isolates typically produce a variety of virions including long filaments (reviewed in (40)), and natural infections seemingly select for filamentous particles (41–43). This suggests that the ability to form filaments might provide an advantage within the host. In support of this idea, it has recently been shown that filamentous virions can enhance infectivity and fusion even in the presence of neutralising antibodies (44). It has also been proposed that the filamentous morphology may enhance direct cell to cell spread of IAV infection (39), which could confer a neutralising antibody evading route of infection spread. We observed that filamentous IAV virions and TLSs are comparable in their dimension, composition, and in the cellular processes involved during their formation (26, 40, 45). For these reasons, we hypothesised that the ability to form filamentous virions might correlate with the ability of IAV to induce TLSs.

To test this, we used A/Puerto Rico/8/1934 H1N1 (PR8), and A/Udorn/307/1972 H3N2 (Udorn) viruses, two strains which are known to retain a characteristic spherical or filamentous virion morphology, respectively (46, 47). By swapping the segment seven of the genome between these viruses, which encodes the matrix protein and with it the primary determinants of virion morphology, we also generated reassortants of Udorn with the matrix of PR8 (Udorn MPR8) and PR8 with the matrix of Udorn (PR8 Mud). These segment seven reassortants display intermediate virion morphologies compared to the wild-type (WT) viruses (48, 49). To measure virion morphology we used immunostaining of haemagglutinin (HA), both on infected MDCK cell membranes during viral budding (S4 Fig), and on free virions that had adsorbed to glass slides (Fig 4A). By quantifying the lengths of filamentous particles within freshly harvested supernatants 48 hours post infection (hpi) we confirmed the predominately spherical and filamentous morphologies of PR8 and Udorn, as well as the intermediate morphologies of Udorn (Udorn MPR8) or PR8 (PR8 MUd) (Fig 4B).

**Fig 4.**
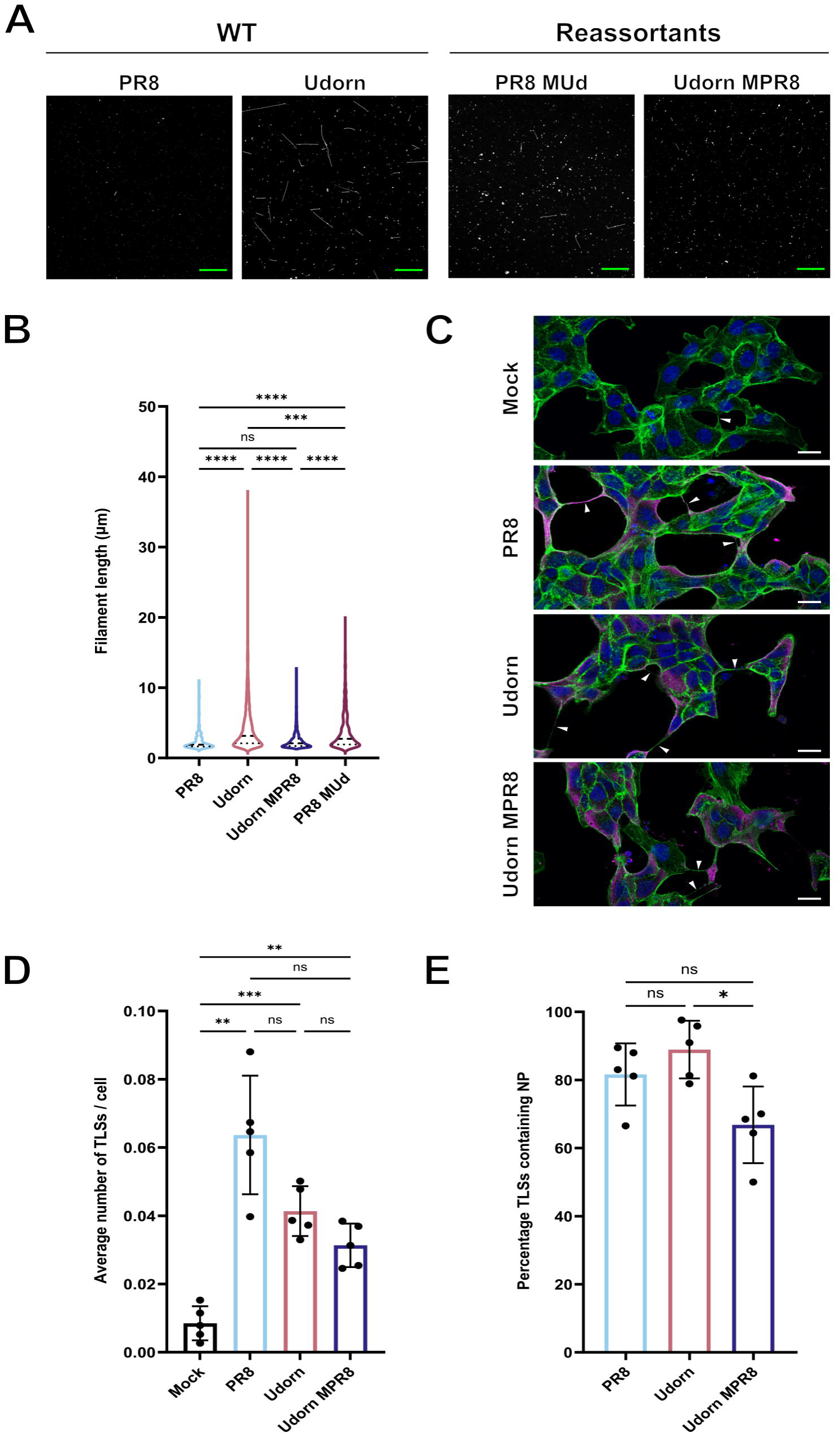
TLS induction is not influenced by IAV virion morphology. **(A)** Representative immunofluorescence images of influenza virions, harvested and fixed on coverslips at 48 h.p.i. and with labelled for hemagglutinin (HA) (white). Scale bars (green) = 20 μm. **(B)** Violin plot of the filament lengths formed by different IAV strains, with the median and the upper and lower quartile values indicated by dashed lines. Individual filament lengths across three independent experiments were plotted. Differences in the mean filament length between strains was tested for significance by Kruskal-Wallis test (n.s. *p* > 0.05, \*\*\**p* < 0.001, **** *p* < 0.0001). **(C)** Representative confocal images of MDCK cells at 16 h after mock, PR8, Udorn or Udorn MPR8 infection. White arrowheads indicate the presence of TLSs. Nuclei (blue), F-actin (green), NP (magenta). Scale bars = 20 μm. For each virus, confocal images of infected cells were used to determine **(D)** the average number of TLSs per cell and **(E)** the percentage of TLSs containing NP. Differences between strains in TLS induction and in the incorporation of NP were tested for significance by one-way ANOVA (n.s. *p* > 0.05, \**p* < 0.05, \*\**p* <0.01, \*\*\**p* < 0.001). The mean and SD from 5 independent experiments is shown.

We then investigated whether the different IAVs induced TLS formation to different extents. To do this, we first infected sub-confluent MDCKs with wild type (WT) PR8 or Udorn at an MOI of 1.5 PFU/cell. At 16 h post infection the cells were fixed, stained for viral NP and F-actin, and the presence of TLSs was quantified using super resolution confocal microscopy (Fig 4C). We found that both viruses induced the formation of TLSs when compared to mock infected cells, and that these TLSs contained NP, suggesting that they could transport viral genomes. However, we found no statistically significant difference in the degree of TLS induction between them (Fig 4D). Indeed, the greatest number of TLSs per cell (though not significantly different from the others) was induced by the spherical virus, PR8. This suggests that TLS induction is a general feature of IAV infection and that virion morphology does not affect this. To confirm this, we utilised the spherical Udorn MPR8 virus and found that the levels of TLS induction was not elevated when compared to WT filamentous Udorn.

Next, we investigated whether virion morphology influenced the frequency of IAV direct cell to cell spread. To do this we implemented a microplaque assay, similar to that used previously by Roberts *et al.* (1). In this assay, a confluent monolayer of MDCKs is infected at a low MOI and incubated under conditions inhibitory to virus particle spread. This included the absence of TPCK trypsin (a protease that cleaves the viral receptor binding protein HA, priming the virus for entry), and the inclusion of the neuraminidase inhibitor (NAi), zanamivir, which prevents the release of newly-formed virions from infected cells. Therefore, the spread of infection to neighbouring cells should only result from direct cell to cell spread rather than from the release of cell free virions. Such a scenario means that infection spread to neighbouring cells causes individual infected cells to develop into NP-positive clusters of adjacent cells, or microplaques (Fig 5A).

**Fig 5.**
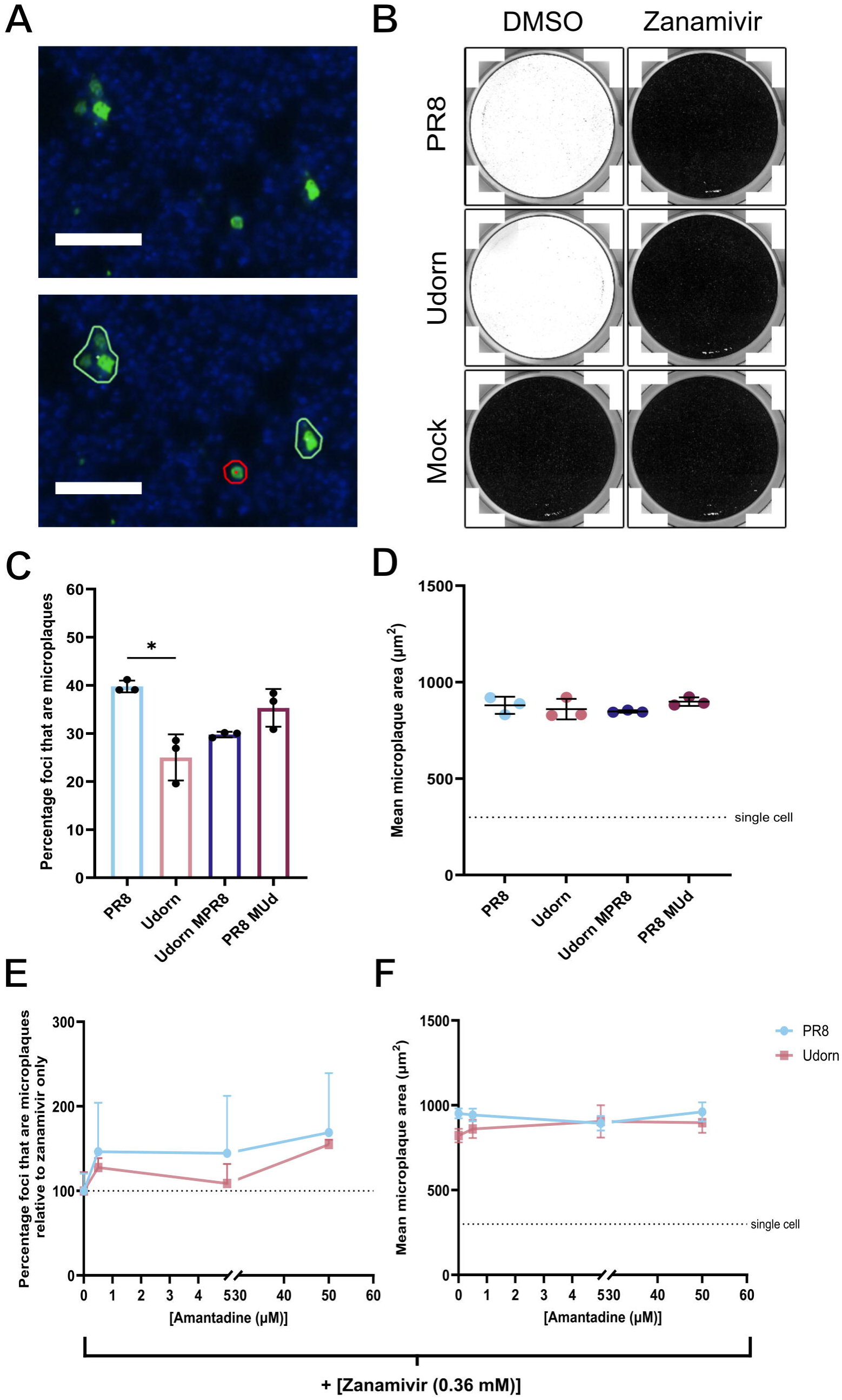
The direct cell-to-cell spread of IAV is independent of virion morphology. **(A)** Representative images of infected MDCKs at 48 h after PR8 infection under microplaque assay conditions, imaged by a Nexcelom Celigo imaging cytometer. Gating thresholds were applied (bottom panel) identifying microplaques (adjacent NP positive cells, circled in green) or isolated infected cells (circled in red). Nuclei (blue), NP (green). Scale bars = 100 µm. **(B)** Coomassie stained MDCK cells 72 h after inoculation with TPCK trypsin treated supernatants, collected from PR8 or Udorn infected microplaque assays performed in the presence of 0.36 mM zanamivir. After absorption, the inoculum was discarded and residual zanamivir was removed by washing with PBS, after which the cells were cultured under an Avicel overlay containing 1 μg/ml TPCK trypsin. Images are representative of two biological repeats. MDCK cells were infected with WT or segment 7 reassortant viruses in the presence of 0.36 mM zanamivir for 48 h and **(C)** the percentage of NP positive foci that are microplaques and **(D)** the mean microplaque area were determined. The significance of differences between strains was determined using a Kruskal-Wallis test (n.s. *p* > 0.05, * p < 0.05). MDCK cells were infected with WT PR8 or Udorn virus in the presence of 0.36 mM zanamivir and varying concentrations of amantadine for 48 h. **(E)** The proportion of NP positive foci that are microplaques, relative to zanamivir only treatment, and **(F)** the mean microplaque area are shown. Differences between strains at each concentration were tested for significance by a Mann-Whitney test, and differences between the same strain at different concentrations were tested by Kruskal-Wallis test (n.s. *p* > 0.05). For all data the mean and SD are shown (n = 3).

We demonstrate that TPCK trypsin is required for PR8 and Udorn to cause cytopathic effect (CPE) and that zanamivir (even at low concentrations) was sufficient to prevent CPE even in the presence of TPCK trypsin (S5A Fig). This suggested that the conditions of this assay was inhibitory to virion spread. To confirm that the production of infectious virions was being inhibited by zanamivir, we excluded TPCK trypsin and infected cells with PR8 and Udorn in the presence or absence of 0.36 mM zanamivir. We then collected supernatants 48 hpi. and incubated them with TPCK trypsin to cleave and activate any released virions. We then performed a plaque assay on new MDCK cells to detect any infectious virus, rinsing cells after absorption to remove residual zanamivir and adding TPCK trypsin to the agarose overlay. While near-complete cell loss occurred when the plaque assay was set up with media from cells infected without zanamivir, no plaques were seen if the media was collected from cells infected in the presence of 0.36 mM zanamivir. Overall, this demonstrates that excluding exogenous proteases prevents multicycle replication at the point of viral entry, but this still results in the production of progeny virus that is capable of infection. This is further supported by the observation that in the absence of TPCK trypsin, the inclusion of zanamivir, even at concentrations less than 0.36 mM, results in an immediate reduction in the frequency and scale of microplaques (S5B & D Fig), with both viruses having similar sensitivities to zanamivir treatment (S5C Fig). Therefore, in order to prevent any spread of infectious cell free virus particles, we included zanamivir in our experimental design.

Having established conditions that robustly prevented the release of infectious virions, we compared the frequency of direct cell to cell spread of IAVs of differing morphologies by quantifying the size and number of microplaques that had formed 48 h post-infection. We found that the IAVs were all capable of direct cell to cell spread, with more than 25% of infected cells infecting their neighbours even when in the absence of virus particle release. The ability to form filamentous virions did not provide any advantage in cell to cell spread (Fig 5C). Indeed the only difference between the strains was a slight increase in the proportion of cells infecting their neighbours for the spherical strain PR8 when compared to the filamentous strain Udorn (*p =* 0.0134, Kruskal-Wallis test), and as this difference was not recapitulated when the matrix gene segment was swapped between these viruses it is probable that this difference was independent of morphology (Fig 5C). Furthermore, in cases where microplaques formed, their eventual size was not influenced by virion morphology (Fig 5D). Together, this indicates that the ability of IAVs to undergo direct cell to cell spread is not determined by virion morphology.

We hypothesised that the difference in the efficiency of microplaque formation between PR8 and Udorn (Fig 5C) could result from a strain dependent preference for the mechanism of direct cell to cell spread. Currently, two parallel mechanisms for the direct cell to cell spread of IAVs have been proposed, which could operate simultaneously: the transfer of cell associated viruses, requiring entry into the endosomal pathway following internalisation of virus particles (2), and the transfer of viral genomes directly between cells with involvement of the actin cytoskeleton (1). To distinguish these, we utilised amantadine (an M2 ion channel blocker which prevents virion uncoating during entry) to assess whether the IAV strains we used differed in their reliance on each of these mechanisms for cell to cell transmission.

We first characterised the antiviral effect of amantadine in a microplaque assay (S6 Fig). We confirmed previous reports that PR8 and Udorn have some resistance to amantadine in the presence of TPCK trypsin, however, amantadine does reduce CPE in a concentration dependent fashion (S6 Fig). Once virus particle release is completely inhibited by zanamivir, increasing concentrations of amantadine caused no further reductions in microplaque formation and area (Fig 5E and F). This suggests that the transfer of cell associated viruses, which should be inhibited by amantadine, cannot be responsible for the high efficiency of direct cell to cell spread of PR8 and Udorn infection that we observed in the presence of amantadine and zanamivir and in the absence of TPCK trypsin (Fig 5C). The alternative is a mechanism that can transfer cytoplasmic viral genomes directly from cell to cell, such as trafficking through open-ended TLSs. Our observation that PR8 and Udorn induce TLSs and incorporate NP at similar frequencies (Fig 4D & E) would suggest that other mechanisms of viral genome transfer are responsible for the differences in the frequency of direct cell to cell spread observed between these viruses (Fig 5C).

### IAVs induce TNT-like structures by the triggering of apoptosis

Thus far our data revealed that TLS induction is a common feature of IAV infection and that the mechanism of induction does not involve extracellular signals, is U.V. sensitive and requires replicating virus. This suggested that the induction of TLSs by IAVs follows an intracellular virus-host interaction. One of the most striking responses of the cell to IAV infection is the onset of apoptosis, and previous reports have found that apoptosis can trigger the formation of TLSs between stressed and healthy cells (50, 51). U.V. inactivated IAVs fail to induce apoptosis (52), and as is often the case in IAV infections we observed significant evidence of apoptosis (nuclear fragmentation, membrane blebbing and apoptotic bodies) under infection conditions where TLSs were induced (Fig 3B iii, 3G and 4C). Therefore, we hypothesised that the triggering of apoptosis by IAVs is required for the induction of TLSs.

To test this, we first used the pan-caspase inhibitor Z-VAD-fmk at increasing concentrations in MDCK cells infected with BrightFlu, a modified PR8 virus encoding the fluorophore ZsGreen (53), at an MOI of 1.5 PFU/cell. At 16 h post infection, apoptotic (positive for active caspase 3/7) and infected (ZsGreen positive) cells were detected using an imaging cytometer, and cell populations were classified by FlowJo analysis (Fig 6A). We found that infection resulted in approximately 95% of cells being positive for ZsGreen signal, with 35% also positive for active caspase 3/7 (Fig 6B). Addition of increasing concentrations of Z-VAD-fmk (up to 100 µM) reduced the percentage of infected cells that were positive for active caspase 3/7 from approximately 35% to 16% (Fig 6B). Importantly, we found that Z-VAD-fmk demonstrated no antiviral effect with no reduction in the percentage of cells infected (∼96% of cells ZsGreen positive).

**Fig 6.**
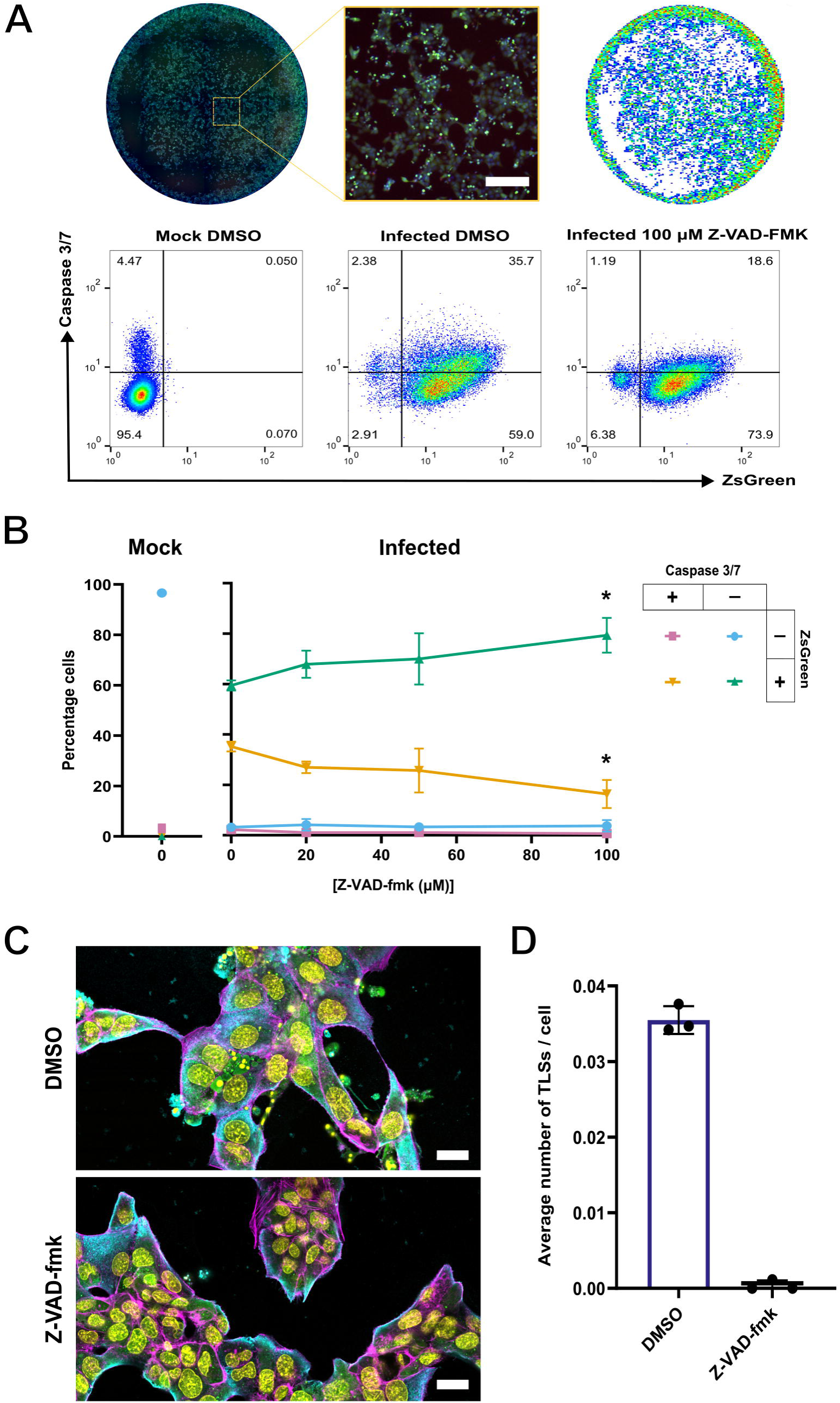
The pan-caspase inhibitor Z-VAD-fmk rescues infected cells from apoptosis and prevents the induction of TLSs. **(A)** Representative images of a scanned well, alongside a magnified inset (nuclei stain (blue), BrightFlu ZsGreen (green), active Caspase 3/7 (red). Scale bar = 200 μm) and nuclei masks following imaging with Nexcelom celigo image cytometer 16 h.p.i (upper panel). Representative FlowJo analyses plots (lower panel) of mock or PR8-ZsGreen (BrightFlu) infected MDCK cells, treated with either DMSO (mock) or Z-VAD-fmk. Cell populations were classified according to the expression of ZsGreen as a marker of infection, and the presence of active Caspase 3/7 as a marker of apoptosis. Gating’s were established according to a Mock DMSO control in the absence of active Caspase 3/7 detection reagent. **(B)** The percentage of MDCK cells positive or negative for active Caspase 3/7 and ZsGreen, at 16 h.p.i. with BrightFlu at an MOI of 1.5 PFU/cell, in the presence of DMSO (mock) or of increasing concentrations of Z-VAD-fmk. The reductions in the percentage of cells singly positive for ZsGreen, or doubly positive for caspase 3/7 and ZsGreen, when compared to the infected DMSO control, were tested for significance using a Kruskal-Wallis test (n.s. *p* > 0.05, \**p* < 0.05). **(C)** Representative confocal images of BrightFlu infected MDCK cells, treated with DMSO or 100 µM Z-VAD-fmk. Nuclei (yellow), F-actin (magenta), ZsGreen (green), NP (blue). Scale bar = 20 µm. **(D)** Average number of TLSs per MDCK cell at 16 h.p.i. with BrightFlu at an MOI of 1.5 PFU/cell in the presence of DMSO or 100 µM Z-VAD-fmk. For all data the mean and SD are shown (n = 3).

We then applied 100 µM Z-VAD-fmk to BrightFlu infected (MOI 1.5 PFU/cell) sub-confluent MDCKs and quantified the TLSs that formed. BrightFlu infection led to an increase in classical cytological indicators of apoptosis, such as nuclear fragmentation and apoptotic bodies (Fig 6C), and as before correlated with an induction of TLSs (Fig 6D). However, upon addition of Z-VAD-fmk there were no visible signs of apoptosis (Fig 6C), and the induction of TLSs also ceased (Fig 6D). We therefore concluded that the induction of TLSs by IAV infection results from the onset of virus-induced apoptosis.

## Discussion

The formation of TLSs has been shown to mediate the direct cell to cell spread of IAV infections in *in vitro* studies (5). However, it was unknown if this effect could occur in the crowded, mobile environment of the respiratory epithelium, or how IAV infection induced the formation of TLSs.

Here, we provided the first evidence that TLSs do form at the sites of IAV infections, using an *in vivo* reporter mouse system to demonstrate that TLSs can form from IAV infected cells in the airway epithelium. This shows that TLSs have the potential to contribute to the within-host spread of IAV infections and also expands the range of tissues known to be conducive to TLS growth (19).

We then used *in vitro* assays to identify the factors behind TLS induction. We found that the induction of TLSs is driven by the presence of replicating virus. This effect was not mediated through paracrine signalling between cells or through IAV’s ability to form filamentous virions that resemble TLSs. Instead, IAV induces TLS formation through triggering apoptosis (Fig 6). Apoptosis was already known to contribute to the pathology of IAV infection (54), and our study shows that it can also be used by the virus to establish routes for direct cell to cell spread.

Our data shows no evidence that IAV infection establishes the conditions for TLS pathfinding (Fig 2), and we show that in a co-culture, uninfected cells produce TLSs at a similar frequency to IAV infected cells (Fig 2B), in line with previous studies (3, 30, 65). Because of this, the contribution of TLSs to IAV infection spread is likely driven by the local density of infected cells at any given point in time. For example, early in an infection where most cells in a region are uninfected, TLS formation would primarily result connections forming between infected and uninfected cells, and so could increase the likelihood of infecting new cells. Later in an infection, when more cells are infected, TLSs between two infected cells might be able to mediate coinfection. Coinfection of a single cell can only happen if entry of additional IAVs occurs during the early stages of the initial infection, due to the time-dependent onset of superinfection exclusion (66), and the rapid delivery of viral genomes by TLSs could help to overcome this (67). Furthermore, a greater number of viral genomes entering a cell can result in complementation between genomes and more rapid viral replication, so the formation of TLSs might indirectly support the production of more cell-free virions (68). In this way TLSs may influence the within-host kinetics of virus production, lesion expansion and possible lesion interactions.

Apoptosis can be distinguished from other cell-death programmes such as necroptosis and pyroptosis by its reliance on caspases, and by the limited involvement of extracellular pathogen- or damage associated molecular patterns, respectively, as stimuli (reviewed in (55, 56)). Recent studies have demonstrated that caspase 8 is also involved in the activation of pyroptosis, in a process independent of extracellular PAMPS or TNF (reviewed in (56)). However, there are redundant pathways for pyroptosis activation, with nuclear programmed-death ligand 1 (nPD-L1) also capable of progressing the cascade (57). Furthermore, treatment of IAV infected dendritic cells and mouse embryonic fibroblasts with a pan-caspase inhibitor did not prevent necroptosis (58, 59). Therefore, the pan-caspase inhibitor Z-VAD-fmk can be considered primarily as an inhibitor of apoptosis in our experiments, in line with its use in other studies (50, 60).

How apoptosis induces TLS formation is complex, as multiple stages of apoptosis can be important for TLS induction. For example, one study found that mitochondrial cytochrome c release within U.V. treated PC12 cells induced the formation of microtubule-containing TLSs (50). Another study found that the presentation of phosphatidylserine (PS) on the outer leaflet of the plasma membrane was required (51). Coating PS with annexin V prevented TLS induction from stressed cells but had no effect on the frequency of other cellular projections (51). This suggests that the presentation of PS on the extracellular surface could increase the rate at which cell projections can interact with recipient cell membranes thereby inducing the formation of TLSs. This could also extend to the observation that uninfected cells were capable of initiating TLS connections under low MOI conditions (Fig 2G). We hypothesise that under such conditions the presentation of PS from infected cells could mediate more intercellular membrane interactions with cell projections originating from uninfected cells. This is supported by the finding that there is a greater likelihood for TLS connections to involve at least one infected cell (approximately 60% of TLS connections involved at least one infected cell despite infection of only 37% of cells, S3 Fig.).Therefore, by acting upstream of these steps in the apoptotic pathway, IAV may be able to influence TLS induction in multiple ways.

Apoptosis is a cell death programme which is already known to influence IAV virus titres *in* vitro and can be seen *in vivo* following IAV infection (52, 61, 62). IAVs have been shown to orchestrate apoptosis in ways that benefit virus propagation (52, 63). The increased formation of TLSs following the onset of apoptosis could also benefit the virus by facilitating the direct cell to cell spread of IAV genomes, and so spreading IAV to a healthy new host cell that can continue viral replication. There may also be other advantages to the virus – for example, it has been shown that mitochondrial trafficking through TLSs can rescue porcine reproductive and respiratory syndrome virus infected cells from death by apoptosis (50, 51, 64). It would be interesting to see if IAV infection also promotes intercellular exchange of mitochondria, and if this could delay cell death and prolong IAV replication within the primary infected cell.

For some other viruses viral proteins are known to be involved in the induction of TLSs (reviewed in (11)). For example, the Nef and Env protein of HIV-1, as well as the US3 proteins of the alphaherpesviruses pseudorabies virus (PRV), Herpes-Simplex virus 2 (HSV-2), and bovine herpesvirus 5 (BoHV-5), have been shown to trigger TLS formation (65, 69–72). We note that these proteins are also known to regulate apoptosis (70, 72–75). Interestingly, in the case of HIV-1 Nef and the US3 proteins of alphaherpesviruses, the ability to induce TLSs correlated to their function in protecting cells from apoptosis (72, 73, 76, 77), with incidences of their ability to form complexes with known TNT forming actin modulating proteins such as phosphoinositide 3-kinase (PI3K) (73). The relationship between virus-induced apoptosis and TLS induction is understudied, and further study of the role of IAV proteins in TLS formation and the regulation of apoptosis may be relevant to our understanding of multiple viruses.

In this study, we reveal how IAVs can establish direct cell-to-cell spread and seed new infections, even in the presence of effective antiviral drugs. We report for the first time structures resembling TNTs within the lungs of infected mice, suggesting that TLS-mediated spread of infections could occur within the respiratory epithelium, and we show that IAV induces TLS formation by driving infected cells into apoptosis. These data suggest that virus-induced apoptosis plays a previously unappreciated role in the spread of IAV, and potentially in the spread of many other viruses, within the tissues of their infected hosts.

## Material & Methods

### Cells, plasmids and viruses

Madin-Darby Canine Kidney (MDCK) cells, human embryonic kidney 293T cells (HEK293T), and adenocarcinoma 549 (A549) cells were maintained in Dulbecco’s Modified Eagle Medium (DMEM, Gibco) supplemented with 10% Foetal Bovine Serum (FBS, Gibco). Cells were maintained at 37°C and 5% CO_2_ in a humidified incubator.

A549 AcGFP1 cells were generated using prepackaged lentivirus (TakaraBio, rLV.EF1.AcGFP1-Mem-9). Transduction of A549 cells was performed as per the manufacturer’s instructions. Briefly, A549s were seeded into a 6 well plate at a density of 2.5X10^5^ cells per well. Transduction mix was prepared by adding lentivirus to complete media containing polybrene (4 μg/ml) to achieve an MOI of 10. The transduction mix was added directly over cells and incubated for 5.5 h at 37°C. The cells were washed once with PBS and complete media added. Following 48 hours incubation the cells were treated and maintained with complete media supplemented with 2 μg/ml puromycin (Thermo) selecting for transduced cells.

Wild-type A/Puerto Rico/8/1934 H1N1 (PR8), and A/Udorn/307/1972 H3N2 (Udorn) P0 viruses were generated as previously described (78). Briefly, HEK293T cells were transfected with the 8 plasmid pDUAL (kindly provided by Prof. Ron Fouchier, Erasmus MC) and 12 plasmid pHH21 (kindly provided by Prof. Paul Digard, Roslin) reverse genetic systems, respectively, and propagated on MDCK cells. Segment 7 reassortant viruses containing the matrix gene of PR8 or Udorn within a background of Udorn (Udorn MPR8) or PR8 (PR8 MUd), were prepared by swapping the corresponding vRNA encoding plasmids. BrightFlu (PR8 virus encoding the fluorophore ZsGreen in segment 8 of the genome, (53)), and PR8-Cre (PR8 virus encoding the Cre recombinase, kindly provided by Professor Ben tenOever) stocks were propagated on MDCK cells. The propagation of all viruses was performed in viral growth media (VGM) (serum-free DMEM supplemented with 1 μg/ml TPCK-treated Trypsin and 0.14% Bovine Serum Albumin (BSA) Fraction V, Sigma) except where stated otherwise. For viral stocks, media were collected 48 h post infection and cell debris removed by centrifugation at 3000 rpm for 5 minutes; aliquots were stored at -70°C. Infectious titres of viruses (as plaque forming units (PFU)/mL) were determined under agarose by conventional plaque assay in MDCKs (79).

### IAV infection of sub-confluent cells

MDCK cells were seeded onto 13 mm glass coverslips at a density of 6×10^4^ cells. Following overnight incubation cells were washed with PBS and infected with WT or reassortant virus at an MOI of 1.5 PFU/cell. Adsorption was performed for 45 minutes AT 37°C in VGM, after which the inoculum was removed, and cells washed with PBS to remove any remaining TPCK trypsin. The cells were overlaid with 1 ml Opti-MEM (Gibco) and sixteen hours post infection formaldehyde was added directly to the Opti-MEM overlay to a final concentration of 4%. Fixation was performed at 8°C for 2 h, and coverslips were then allowed to air dry for approximately 10 minutes before washing once with 2% (v/v) FBS/PBS.

For the A549 and A549 AcGFP1 coculture, cells were seeded at a 1:1 ratio to a density of 4.5×10^4^ cells. Following overnight incubation, cells were infected with PR8 at an MOI of 2.5 PFU/cell with the overlay conditions and fixation method the same as previously stated.

### Mice infections and thick tissue sectioning

Animal work was done in accordance with the EU Directive 2010/63/eu and Animal (Scientific Procedures) Act 1986, under a project licence P72BA642F. Three mT/mG mice, between 17 and 18 weeks old, were kindly provided by Dr Stephanie May (Cancer Research UK, Scotland Institute). Two mice were intranasally infected with 1000 PFU of PR8-Cre. A naïve mouse was mock infected with PBS in a similar manner. Six days post infection the mice were euthanised, the lungs harvested and inflated with agarose. Lungs were fixed in 4% formaldehyde overnight before being transferred to PBA (1% BSA, 0.05% NaN_3_ in PBS). Dissected lung lobes were embedded in optimal cutting temperature compound (OCT, VWR) with 100 µm tissue sections cut at -20°C using a CryoStat (CM1950, Leica) and slides stored at -70°C. Prior to imaging, samples were thawed, and OCT removed by immersing in PBS. Coverslips were placed on seal-frame incubation chambers (Thermo Fisher) containing Ce3D tissue clearing solution.

### Immunocytochemistry and immunostaining of virus particles

For virion staining, MDCKs were infected with P1 virus at an MOI of 0.25 PFU/cell. 48 h post infection the supernatant was collected, and cell debris removed by centrifugation at 13,000 rpm for 1 minute. The virus supernatant was diluted 1:10 in PBS and spun onto coverslips by centrifugation at 1000 *g* for 30 minutes at 4°C. Coverslips were fixed for 10 minutes with 4% formaldehyde and blocked in 2% FBS/PBS. For all viruses with the exception of PR8 MUd, coverslips were stained with mouse anti-HA primary antibody [1:2000] (kindly provided by Dr Stephen Wharton, Francis Crick Institute) and goat anti-mouse Alexa-Fluor 555 (Thermo Fisher). PR8 MUd was stained using rabbit anti-H1 (kindly provided by Prof. Paul Digard, University of Edinburgh, (80)) [1:100]. For HA surface-stained cells, unpermeabilised cells were blocked with 2% FBS/PBS for one hour with immunostaining performed as above with the inclusion of 4′,6-diamidino-2-phenylindole (DAPI, Thermo Fisher) [1:1000].

To achieve intracellular staining, cells were permeabilised with 0.2% Triton-X100 in PBS for 7 minutes and washed three times with 2% FBS/PBS. Samples were blocked in 2% FBS/PBS for 1 h followed by the addition of a sheep anti-NP antibody (PR8 H1N1 available from the Influenza Virus Toolkit at www.influenza.bio) [1:1000] and incubated for 1 h at room temperature. Cells were washed three times before secondary antibody incubation for 30 minutes using anti-sheep alexa fluor 488, 555 or 647 (Thermo Fisher) alongside DAPI [1:1000]. For super resolution confocal microscopy, cells were additionally stained with 1X phalloidin-iFluor 488, 555 or 647 (abcam) in 1% bovine serum albumin (BSA) for 30 minutes. Coverslips were mounted onto slides using ProLong Gold Antifade mounting media.

### Microplaque Assay

Confluent MDCK cells were infected with PR8, Udorn or Udorn MPR8 at MOIs achieving approximately 1000 nucleoprotein (NP) positive infected foci per well. Following a 2 h incubation, the inoculum was removed, and cells acid washed (PBS-HCl pH 3) for 1 minute to inactivate free virions. Cells were washed three time with PBS before adding 1 ml overlay containing 1.2% Avicel, DMEM, 0.14% BSA and between 0.12 to 0.48 mM zanamivir (Sigma). Where indicated, amantadine (1-adamantanamine, Thermo Fisher) was included at concentrations ranging from 0.5 mM to 50 mM in place, or in combination with 0.36 mM Zanamivir.

Forty-eight h post infection, the overlay was removed by washing with PBS. Cells were fixed with either 4% formaldehyde for 10 minutes or Coomassie Blue for 30 minutes.

### U.V. inactivation of virus and western blotting

Supernatants were collected and distributed in 100 µl volumes in a 96-well plate. Inactivation of virus was performed using an 8W 254 nm U.V. lamp (UVS-28 EL, UVP) placed directly above samples whilst on ice for 6 minutes, with shaking every 2 minutes. U.V. treated samples were collected and added directly to fresh cells.

At specified time points cells were harvested and lysed in Laemmli buffer (20% glycerol, 2% SDS, 24 mM Tris pH 6.8, 0.1M DTT, 0.2% bromophenol blue, 0.2% cyanol) supplemented with benzonase (Merck) and protease inhibitor cocktail (Thermo Fisher). Samples were heated at 37°C for 30 minutes, then at 95°C for 5 minutes. Samples were run on AnyKD mini-PROTEAN TGX gels (BioRad) before being transferred to a nitrocellulose membrane using the TransBlot Turbo quick transfer protocol. Membranes were blocked for 1 h with 0.1% Tween 20 (PBS-T)/5% milk, washed three times with PBS-T and stained overnight at 4°C in PBS-T/5% milk containing primary antibodies (anti-phosphorylated STAT1, Cell Signalling Technology and anti-alpha Tubulin, Merck) [1:1000]. The membranes were washed three times with PBS-T before secondary antibody incubation with anti-rabbit DyLight 800 and anti-Mouse DyLight 680 (Invitrogen) diluted in PBS-T/5% milk [1:10,000] for 45 minutes. Membranes were washed three times with PBS-T, PBS and water prior to imaging using the LI-COR CLx-Odyssey Imaging platform with quantification performed using the Image Studio Lite Software.

### Detection of active caspase 3/7 and inhibition of apoptosis

MDCKs were infected at an MOI of 1.5 PFU/cell with BrightFlu. Following virus absorption, an overlay consisting of DMEM, 60 μM caspase 3/7 red detection reagent (Thermo Fisher), and the pan-caspase inhibitor Z-VAD-fmk (Promega) were added to the cells and incubated for 16 h. Hoescht 33342 was added directly to the overlay achieving a final concentration of 5 µg/mL and incubated for 20 minutes. Cells were imaged live using the Nexcelom Celigo image cytometer.

### Confocal microscopy

Confocal microscopy was performed using the Zeiss LSM 880 (63x oil immersion objective, 1.4 NA). Super resolution imaging of TLSs and budding filaments was performed using Airyscan fast detection. Post-acquisition auto processing was performed within Zen Black (Zeiss) software (v14.0.29.201). For TLS quantification, a single field of view encompassed two adjacent tiles stitched together. In total 14 randomly selected fields of view with a suitable distribution of cell nuclei were selected per technical replicate. Each biological replicate consisted of two technical replicates. Imaging of filaments on coverslips was performed similarly except for the sole use of the Gallium Arsenide Phosphide (GaAsP) detector.

Thick section confocal microscopy was similarly performed utilising Z-stacks encompassing 3D regions of interest. Maximum intensity projections of 3D airyscan processed Z-stacked images were created using the Zen Black (Zeiss) software (v14.0.29.201).

### Image analysis

Three-dimensional surface renders of fluorescent objects within thick tissue sections were created from Z-stack images via the creation of binary masks on Imaris (Andor). Any further background removal required was performed within Imaris according to samples derived from naïve mice.

Filament measuring was performed using custom image J macro scripts published previously (81). Briefly, micrographs were auto thresholded, and debris with a circularity between 0.5 and 1 was removed via Particle Remover. The dimensions of quantified remaining particles were extracted with Ridge Detection and figures created on GraphPad prism.

TNT-like structures (TLS) were quantified manually using characteristic features of TNTs that differentiate them from other protrusions. These include the presence of a narrow structure that appears to connect two or more cells and contains F-actin along its length. The structure is forming from cells that have a visibly intact nucleus and are not showing signs of cell division. Additionally, structures having cellular midbodies were excluded as having likely resulted from recent cell division as have been done in previous analysis (30). For A549s a length threshold of 5 μm was implemented to distinguish TLS from filopodia that we observed to be more common in this cell type. Quantification of micrographs was performed blind to avoid analyst bias.

Microplaque image analysis was performed on the Nexcelom Celigo image cytometer using a 90% well mask. A gating area of 600 μm^2^ was selected on the Celigo analysis software to distinguish microplaques (i.e. areas with Alexa fluor 488 signal > 600 μm^2^) from single infected foci. The percentage of total foci that existed as microplaques was determined alongside the mean microplaque area as measured by the Celigo.

The detection of cells caspase 3/7 and ZsGreen positive was performed on a Celigo imaging cytometer. A nuclei mask on the Hoechst channel with high precision within the analysis software on the Celigo imaging cytometer. A dilation radius of 5 μm was applied to capture perinuclear and cytoplasmic red or green signal. The percentage of cells negative, singly positive, or dual positive for active caspase 3/7 and ZsGreen was performed within FlowJo software (v10.6). Cell population gating was established based on mock infected nuclei-stained controls and then applied to all samples.

## Abbreviations

BSA: bovine serum albumin
CPE: cytopathic effect
DMEM: Dulbecco’s Modified Eagle Medium
GFP: green fluorescence protein
HIV, HA: Hemagglutinin; human immunodeficiency virus
HSV: herpes simplex virus
HTLV: human T-lymphotropic virus
IFN: interferon
IAV: influenza A virus
MDCK: Madin–Darby canine kidney
MOI: multiplicity of infection
NAi: neuraminidase inhibitor
NP: nucleoprotein
PFU: plaque-forming unit
PRV: pseudorabies virus
PS: phosphatidylserine
SARS-CoV-2: Severe Acute Respiratory Syndrome Coronavirus 2
TNTs: tunnelling nanotubes
TLS: tunnelling nanotube-like structures
U.V.: ultraviolet light, VGM, viral growth media
WT: wild-type

## Funding

This work was supported by funding from the UK Medical Research Council (MRC), as a studentship to D.W [MC_ST_00034] and as core funding to C.L [MC_UU_12016/10] and Quinquennial funding to the MRC-University of Glasgow Centre for Virus Research [MC_UU_12014/9 and MC_UU_00034/1]. E.H. was funded by a Transition Support Award from 332 the UK Medical Research Council [MR/V035789/1]. We also acknowledge funding the Wellcome Trust as Four-Year PhD studentships in Basic Science 325 to C.B.A [226861/Z/23/Z], and J.M [218518/Z/19/Z]. This work was also supported by Cancer Research UK (CRUK) core funding to the Scotland Institute [A31287] and to E.R. [A1920]. The funders had no role in study design, data collection and analysis, decision to publish, or preparation of the manuscript.

## Acknowledgements

We would like to thank Colin Nixon for performing the Cryostat tissue sectioning as well as Anna Sims for providing advice, assistance in choice of analysis and helpful discussions. To Emma Davies and Matt Turnbull we thank for their advice and the provision of reagents. We would also like to thank Alma Macdnonald for providing access to the U.V. lamp.

## Author contributions

D.W.: conceptualisation, methodology, investigation, visualisation, writing – original draft, writing – review and editing, C.B-A.: methodology, investigation, J.M.: investigation, C.L.: methodology, visualisation, E.R.: methodology, investigation, funding acquisition, E.H.: conceptualisation, funding acquisition, methodology, supervision, writing – review and editing.

## Supplementary Captions

**S1 Fig. A TNT-like structure connecting IAV infected epithelial cells within the lungs of a reporter mouse.** Maximum intensity projection of a TNT-like structure (TLS) within thick sectioned lung tissue connecting IAV infected cells. Magnified inset outlined in a white box is shown alongside a 3D render (bottom right panel). Cell membranes are labelled in tdTomato (magenta) for uninfected cells, and GFP (green) for infected cells. White arrow indicates the presence of a TLS. Scale bars = 20 µm.

**S2 Fig. A549 AcGFP cells demonstrate similar properties to WT A549 in infected coculture.** A549 AcGFP1 cells were cocultured with WT A549 cells at a ratio of 1:1 and infected at a low MOI. 16 h.p.i the cells were fixed and immunostained for NP. Following manual assessment of micrographs, the percentage of cells in coculture that were GFP positive (i.e. A549 AcGFP), GFP positive and infected and the percentage of TLS consisting of GFP labelled membrane throughout its length was determined. The means and standard deviations of three biological replicates are shown.

**S3 Fig. TLSs connect cells with both symmetric and asymmetric infection.** Percentage of TLSs connecting WT and AcGFP labelled A549 cells with both asymmetric and symmetric infection status. The means and standard deviations of 3 biological replicates are shown.

**S4 Fig. Morphology of budding IAV virions.** Maximum intensity projections of surface HA labelled MDCK cells, at 16 h.p.i. with the IAV strains PR8 and Udorn (WT) or the segment 7 reassortant viruses Udorn MPR8 or PR8 MUd. DAPI (blue), HA (yellow). Scale bars = 20 μm

**S5 Fig. The cytopathic effect and intercellular spread of IAV is affected by TPCK trypsin and Zanamivir. (A)** Coomassie stained MDCK cells in 12 well plates, 48 h.p.i. with the IAV strains PR8 or Udorn under microplaque assay conditions. The influence of Zanamivir (0.12 mM) or TPCK trypsin (1 µg/ml) was assessed by cytopathic effect (CPE). Images are representative of two biological replicates. **(B)** The percentage NP positive foci that form microplaques under increasing zanamivir concentrations. Differences between viruses at each concentration were tested for significance by Mann-Whitney test, and differences between concentrations were tested by Kruskal-Wallis test (n.s. *p* > 0.05). The means and standard deviations of three biological replicates are shown. The same data are shown in **(C)** with each virus normalised to its behaviour in the absence of zanamivir. **(D)** Mean microplaque area under increasing zanamivir concentrations, with a dashed line showing the approximate area of a single cell. The means and standard deviations of three biological replicates are shown. The significance of differences between viruses was determined by Kruskal-Wallis test (n.s. *p* > 0.05).

**S6 Fig. Amantadine has a concentration dependent effect on IAV induced cytopathic effect.** Coomassie stained MDCKs within a 12 well plate, 48 h.p.i. with either PR8 or Udorn, in the presence (upper panel) and absence (lower panel) of TPCK trypsin at increasing amantadine concentrations. Images are representative of two biological replicates.

## Notes

### Competing Interest Statement

The authors have declared no competing interest.

## References

1. Roberts KL, Manicassamy B, Lamb RA. Influenza A virus uses intercellular connections to spread to neighboring cells. J Virol. 2015;89(3):1537–49.

2. Mori K, Haruyama T, Nagata K. Tamiflu-Resistant but HA-Mediated Cell-to-Cell Transmission through Apical Membranes of Cell-Associated Influenza Viruses. PloS one. 2011;6:e28178.

3. Kumar A, Kim JH, Ranjan P, Metcalfe MG, Cao W, Mishina M, et al. Influenza virus exploits tunneling nanotubes for cell-to-cell spread. Scientific Reports. 2017;7(1):40360.

4. Kongsomros S, Manopwisedjaroen S, Chaopreecha J, Wang SF, Borwornpinyo S, Thitithanyanont A. Rapid and Efficient Cell-to-Cell Transmission of Avian Influenza H5N1 Virus in MDCK Cells Is Achieved by Trogocytosis. Pathogens. 2021;10(4).

5. Ganti K, Han J, Manicassamy B, Lowen AC. Rab11a mediates cell-cell spread and reassortment of influenza A virus genomes via tunneling nanotubes. PLoS Pathog. 2021;17(9):e1009321.

6. Cifuentes-Munoz N, El Najjar F, Dutch RE. Viral cell-to-cell spread: Conventional and non-conventional ways. Adv Virus Res. 2020;108:85–125.

7. Martinez MG, Kielian M. Intercellular Extensions Are Induced by the Alphavirus Structural Proteins and Mediate Virus Transmission. PLoS Pathog. 2016;12(12):e1006061.

8. Jolly C, Kashefi K, Hollinshead M, Sattentau QJ. HIV-1 cell to cell transfer across an Env-induced, actin-dependent synapse. J Exp Med. 2004;199(2):283–93.

9. Hübner W, McNerney GP, Chen P, Dale BM, Gordon RE, Chuang FY, et al. Quantitative 3D video microscopy of HIV transfer across T cell virological synapses. Science. 2009;323(5922):1743-7.

10. Ambrosini AE, Enquist LW. Cell-Fusion Events Induced by α-Herpesviruses. Future Virology. 2015;10(2):185–200.

11. Jansens RJJ, Tishchenko A, Favoreel HW. Bridging the Gap: Virus Long-Distance Spread via Tunneling Nanotubes. J Virol. 2020;94(8).

12. Rustom A, Saffrich R, Markovic I, Walther P, Gerdes H-H. Nanotubular highways for intercellular organelle transport. Science. 2004;303(5660):1007–10.

13. Austefjord MW, Gerdes HH, Wang X. Tunneling nanotubes: Diversity in morphology and structure. Commun Integr Biol. 2014;7(1):e27934.

14. Ady J, Thayanithy V, Mojica K, Wong P, Carson J, Rao P, et al. Tunneling nanotubes: an alternate route for propagation of the bystander effect following oncolytic viral infection. Molecular Therapy - Oncolytics. 2016;3:16029.

15. Eugenin EA, Gaskill PJ, Berman JW. Tunneling nanotubes (TNT) are induced by HIV-infection of macrophages: a potential mechanism for intercellular HIV trafficking. Cell Immunol. 2009;254(2):142–8.

16. Okafo G, Prevedel L, Eugenin E. Tunneling nanotubes (TNT) mediate long-range gap junctional communication: Implications for HIV cell to cell spread. Scientific Reports. 2017;7(1):16660.

17. Pepe A, Pietropaoli S, Vos M, Barba-Spaeth G, Zurzolo C. Tunneling nanotubes provide a route for SARS-CoV-2 spreading. Sci Adv. 2022;8(29):eabo0171.

18. Ady JW, Desir S, Thayanithy V, Vogel RI, Moreira AL, Downey RJ, et al. Intercellular communication in malignant pleural mesothelioma: properties of tunneling nanotubes. Front Physiol. 2014;5:400.

19. Chinnery HR, Pearlman E, McMenamin PG. Cutting edge: Membrane nanotubes in vivo: a feature of MHC class II+ cells in the mouse cornea. J Immunol. 2008;180(9):5779–83.

20. Zhang J-Q, Takahashi A, Gu J-Y, Zhang X, Kyumoto-Nakamura Y, Kukita A, et al. In vitro and in vivo detection of tunneling nanotubes in normal and pathological osteoclastogenesis involving osteoclast fusion. Laboratory Investigation. 2021;101(12):1571–84.

21. Antanavičiūtė I, Rysevaitė K, Liutkevičius V, Marandykina A, Rimkutė L, Sveikatienė R, et al. Long-distance communication between laryngeal carcinoma cells. PLoS One. 2014;9(6):e99196.

22. Thayanithy V, Dickson EL, Steer C, Subramanian S, Lou E. Tumor-stromal cross talk: direct cell-to-cell transfer of oncogenic microRNAs via tunneling nanotubes. Transl Res. 2014;164(5):359–65.

23. Desir S, Dickson EL, Vogel RI, Thayanithy V, Wong P, Teoh D, et al. Tunneling nanotube formation is stimulated by hypoxia in ovarian cancer cells. Oncotarget. 2016;7(28):43150–61.

24. Muzumdar MD, Tasic B, Miyamichi K, Li L, Luo L. A global double-fluorescent Cre reporter mouse. genesis. 2007;45(9):593–605.

25. Panasiuk M, Rychłowski M, Derewońko N, Bieńkowska-Szewczyk K. Tunneling Nanotubes as a Novel Route of Cell-to-Cell Spread of Herpesviruses. Journal of Virology. 2018;92(10):10.1128/jvi.00090-18.

26. Dagar S, Pathak D, Oza HV, Mylavarapu SVS. Tunneling nanotubes and related structures: molecular mechanisms of formation and function. Biochemical Journal. 2021;478(22):3977–98.

27. Sun X, Wang Y, Zhang J, Tu J, Wang XJ, Su XD, et al. Tunneling-nanotube direction determination in neurons and astrocytes. Cell Death Dis. 2012;3(12):e438.

28. Abounit S, Bousset L, Loria F, Zhu S, de Chaumont F, Pieri L, et al. Tunneling nanotubes spread fibrillar α-synuclein by intercellular trafficking of lysosomes. Embo j. 2016;35(19):2120–38.

29. Naphade S, Sharma J, Gaide Chevronnay HP, Shook MA, Yeagy BA, Rocca CJ, et al. Brief reports: Lysosomal cross-correction by hematopoietic stem cell-derived macrophages via tunneling nanotubes. Stem Cells. 2015;33(1):301–9.

30. Omsland M, Pise-Masison C, Fujikawa D, Galli V, Fenizia C, Parks RW, et al. Inhibition of Tunneling Nanotube (TNT) Formation and Human T-cell Leukemia Virus Type 1 (HTLV-1) Transmission by Cytarabine. Scientific Reports. 2018;8(1):11118.

31. Doceul V, Hollinshead M, van der Linden L, Smith GL. Repulsion of superinfecting virions: a mechanism for rapid virus spread. Science. 2010;327(5967):873–6.

32. Julkunen I, Sareneva T, Pirhonen J, Ronni T, Melén K, Matikainen S. Molecular pathogenesis of influenza A virus infection and virus-induced regulation of cytokine gene expression. Cytokine & Growth Factor Reviews. 2001;12(2):171–80.

33. Short KR, Kroeze EJBV, Fouchier RAM, Kuiken T. Pathogenesis of influenza-induced acute respiratory distress syndrome. The Lancet Infectious Diseases. 2014;14(1):57–69.

34. Omsland M, Andresen V, Gullaksen SE, Ayuda-Durán P, Popa M, Hovland R, et al. Tyrosine kinase inhibitors and interferon-α increase tunneling nanotube (TNT) formation and cell adhesion in chronic myeloid leukemia (CML) cell lines. Faseb j. 2020;34(3):3773–91.

35. Patheja P, Sahu K. Macrophage conditioned medium induced cellular network formation in MCF-7 cells through enhanced tunneling nanotube formation and tunneling nanotube mediated release of viable cytoplasmic fragments. Experimental Cell Research. 2017;355(2):182–93.

36. Melwani PK, Balla MMS, Bhamani A, Nandha SR, Checker R, Pandey BN. Macrophage-conditioned medium enhances tunneling nanotube formation in breast cancer cells via PKC, Src, NF-κB, and p38 MAPK signaling. Cell Signal. 2024:111274.

37. Seitz C, Frensing T, Höper D, Kochs G, Reichl U. High yields of influenza A virus in Madin–Darby canine kidney cells are promoted by an insufficient interferon-induced antiviral state. Journal of General Virology. 2010;91(7):1754–63.

38. Mosley VM, Wyckoff RWG. Electron Micrography of the Virus of Influenza. Nature. 1946;157(3983):263-.

39. Vijayakrishnan S, Loney C, Jackson D, Suphamungmee W, Rixon FJ, Bhella D. Cryotomography of budding influenza A virus reveals filaments with diverse morphologies that mostly do not bear a genome at their distal end. PLoS pathogens. 2013;9(6):e1003413.

40. Dadonaite B, Vijayakrishnan S, Fodor E, Bhella D, Hutchinson EC. Filamentous influenza viruses. J Gen Virol. 2016;97(8):1755–64.

41. Seladi-Schulman J, Steel J, Lowen AC. Spherical influenza viruses have a fitness advantage in embryonated eggs, while filament-producing strains are selected in vivo. J Virol. 2013;87(24):13343–53.

42. Choppin PW, Murphy JS, Tamm I. Studies of two kinds of virus particles which comprise influenza A2 virus strains. III. Morphological characteristics: independence to morphological and functional traits. J Exp Med. 1960;112(5):945–52.

43. Chu CM, Dawson IM, Elford WJ. FILAMENTOUS FORMS ASSOCIATED WITH NEWLY ISOLATED INFLUENZA VIRUS. The Lancet. 1949;253(6554):602-3.

44. Li T, Li Z, Deans EE, Mittler E, Liu M, Chandran K, Ivanovic T. The shape of pleomorphic virions determines resistance to cell-entry pressure. Nature Microbiology. 2021;6(5):617–29.

45. Bruce EA, Digard P, Stuart AD. The Rab11 pathway is required for influenza A virus budding and filament formation. J Virol. 2010;84(12):5848–59.

46. Trifkovic S, Gilbertson B, Fairmaid E, Cobbin J, Rockman S, Brown LE. Gene Segment Interactions Can Drive the Emergence of Dominant Yet Suboptimal Gene Constellations During Influenza Virus Reassortment. Front Microbiol. 2021;12:683152.

47. Kolpe A, Arista-Romero M, Schepens B, Pujals S, Saelens X, Albertazzi L. Super-resolution microscopy reveals significant impact of M2e-specific monoclonal antibodies on influenza A virus filament formation at the host cell surface. Scientific Reports. 2019;9(1):4450.

48. Bourmakina SV, García-Sastre A. Reverse genetics studies on the filamentous morphology of influenza A virus. Journal of General Virology. 2003;84(3):517–27.

49. Hutchinson EC, Charles PD, Hester SS, Thomas B, Trudgian D, Martínez-Alonso M, Fodor E. Conserved and host-specific features of influenza virion architecture. Nature Communications. 2014;5(1):4816.

50. Wang X, Gerdes HH. Transfer of mitochondria via tunneling nanotubes rescues apoptotic PC12 cells. Cell Death & Differentiation. 2015;22(7):1181–91.

51. Liu K, Ji K, Guo L, Wu W, Lu H, Shan P, Yan C. Mesenchymal stem cells rescue injured endothelial cells in an in vitro ischemia-reperfusion model via tunneling nanotube like structure-mediated mitochondrial transfer. Microvasc Res. 2014;92:10–8.

52. Wurzer WJ, Planz O, Ehrhardt C, Giner M, Silberzahn T, Pleschka S, Ludwig S. Caspase 3 activation is essential for efficient influenza virus propagation. The EMBO Journal. 2003;22.

53. Pirillo C, Al Khalidi S, Sims A, Devlin R, Zhao H, Pinto R, et al. Cotransfer of antigen and contextual information harmonizes peripheral and lymph node conventional dendritic cell activation. Sci Immunol. 2023;8(85):eadg8249.

54. Uiprasertkul M, Kitphati R, Puthavathana P, Kriwong R, Kongchanagul A, Ungchusak K, et al. Apoptosis and pathogenesis of avian influenza A (H5N1) virus in humans. Emerg Infect Dis. 2007;13(5):708–12.

55. Yuan J, Ofengeim D. A guide to cell death pathways. Nature Reviews Molecular Cell Biology. 2024;25(5):379–95.

56. Wu Y, Zhang J, Yu S, Li Y, Zhu J, Zhang K, Zhang R. Cell pyroptosis in health and inflammatory diseases. Cell Death Discovery. 2022;8(1):191.

57. Hou J, Zhao R, Xia W, Chang CW, You Y, Hsu JM, et al. PD-L1-mediated gasdermin C expression switches apoptosis to pyroptosis in cancer cells and facilitates tumour necrosis. Nat Cell Biol. 2020;22(10):1264–75.

58. Hartmann BM, Albrecht RA, Zaslavsky E, Nudelman G, Pincas H, Marjanovic N, et al. Pandemic H1N1 influenza A viruses suppress immunogenic RIPK3-driven dendritic cell death. Nat Commun. 2017;8(1):1931.

59. Shubina M, Tummers B, Boyd DF, Zhang T, Yin C, Gautam A, et al. Necroptosis restricts influenza A virus as a stand-alone cell death mechanism. J Exp Med. 2020;217(11).

60. Liu B, Meng D, Wei T, Zhang S, Hu Y, Wang M. Apoptosis and pro-inflammatory cytokine response of mast cells induced by influenza A viruses. PLoS One. 2014;9(6):e100109.

61. Mori I, Komatsu T, Takeuchi K, Nakakuki K, Sudo M, Kimura Y. In vivo induction of apoptosis by influenza virus. J Gen Virol. 1995;76 (Pt 11):2869–73.

62. Ito T, Kobayashi Y, Morita T, Horimoto T, Kawaoka Y. Virulent influenza A viruses induce apoptosis in chickens. Virus Res. 2002;84(1-2):27–35.

63. Wurzer WJ, Ehrhardt C, Pleschka S, Berberich-Siebelt F, Wolff T, Walczak H, et al. NF-kappaB-dependent induction of tumor necrosis factor-related apoptosis-inducing ligand (TRAIL) and Fas/FasL is crucial for efficient influenza virus propagation. J Biol Chem. 2004;279(30):30931–7.

64. Guo R, Davis D, Fang Y. Intercellular transfer of mitochondria rescues virus-induced cell death but facilitates cell-to-cell spreading of porcine reproductive and respiratory syndrome virus. Virology. 2018;517:122–34.

65. Sherer NM, Lehmann MJ, Jimenez-Soto LF, Horensavitz C, Pypaert M, Mothes W. Retroviruses can establish filopodial bridges for efficient cell-to-cell transmission. Nature cell biology. 2007;9(3):310–5.

66. Sims A, Tornaletti LB, Jasim S, Pirillo C, Devlin R, Hirst JC, et al. Superinfection exclusion creates spatially distinct influenza virus populations. PLoS Biol. 2023;21(2):e3001941.

67. Tiwari V, Koganti R, Russell G, Sharma A, Shukla D. Role of Tunneling Nanotubes in Viral Infection, Neurodegenerative Disease, and Cancer. Front Immunol. 2021;12:680891.

68. Jacobs NT, Onuoha NO, Antia A, Steel J, Antia R, Lowen AC. Incomplete influenza A virus genomes occur frequently but are readily complemented during localized viral spread. Nat Commun. 2019;10(1):3526.

69. Xu W, Santini PA, Sullivan JS, He B, Shan M, Ball SC, et al. HIV-1 evades virus-specific IgG2 and IgA responses by targeting systemic and intestinal B cells via long-range intercellular conduits. Nat Immunol. 2009;10(9):1008–17.

70. Jansens RJJ, Van den Broeck W, De Pelsmaeker S, Lamote JAS, Van Waesberghe C, Couck L, Favoreel HW. Pseudorabies Virus US3-Induced Tunneling Nanotubes Contain Stabilized Microtubules, Interact with Neighboring Cells via Cadherins, and Allow Intercellular Molecular Communication. J Virol. 2017;91(19).

71. Finnen RL, Roy BB, Zhang H, Banfield BW. Analysis of filamentous process induction and nuclear localization properties of the HSV-2 serine/threonine kinase Us3. Virology. 2010;397(1):23–33.

72. Ladelfa MF, Kotsias F, Del Médico Zajac MP, Van den Broeke C, Favoreel H, Romera SA, Calamante G. Effect of the US3 protein of bovine herpesvirus 5 on the actin cytoskeleton and apoptosis. Vet Microbiol. 2011;153(3-4):361–6.

73. Wolf D, Witte V, Laffert B, Blume K, Stromer E, Trapp S, et al. HIV-1 Nef associated PAK and PI3-kinases stimulate Akt-independent Bad-phosphorylation to induce anti-apoptotic signals. Nat Med. 2001;7(11):1217–24.

74. Perfettini JL, Castedo M, Roumier T, Andreau K, Nardacci R, Piacentini M, Kroemer G. Mechanisms of apoptosis induction by the HIV-1 envelope. Cell Death Differ. 2005;12 Suppl 1:916–23.

75. Murata T, Goshima F, Yamauchi Y, Koshizuka T, Takakuwa H, Nishiyama Y. Herpes simplex virus type 2 US3 blocks apoptosis induced by sorbitol treatment. Microbes and Infection. 2002;4(7):707–12.

76. Wang Y, Cui J, Sun X, Zhang Y. Tunneling-nanotube development in astrocytes depends on p53 activation. Cell Death Differ. 2011;18(4):732–42.

77. Geenen K, Favoreel HW, Olsen L, Enquist LW, Nauwynck HJ. The pseudorabies virus US3 protein kinase possesses anti-apoptotic activity that protects cells from apoptosis during infection and after treatment with sorbitol or staurosporine. Virology. 2005;331(1):144–50.

78. Fodor E, Devenish L, Engelhardt OG, Palese P, Brownlee GG, García-Sastre A. Rescue of influenza A virus from recombinant DNA. J Virol. 1999;73(11):9679–82.

79. Gaush CR, Smith TF. Replication and plaque assay of influenza virus in an established line of canine kidney cells. Appl Microbiol. 1968;16(4):588–94.

80. Amorim M-J, Read EK, Dalton RM, Medcalf L, Digard P. Nuclear Export of Influenza A Virus mRNAs Requires Ongoing RNA Polymerase II Activity. Traffic. 2007;8(1):1–11.

81. Hirst JC, Hutchinson EC. Single-particle measurements of filamentous influenza virions reveal damage induced by freezing. J Gen Virol. 2019;100(12):1631–40.

