## Supplementary figures and images for "Influenza A viruses induce tunnelling nanotube-like structures through the onset of apoptosis"

### Supplementary Figure 1

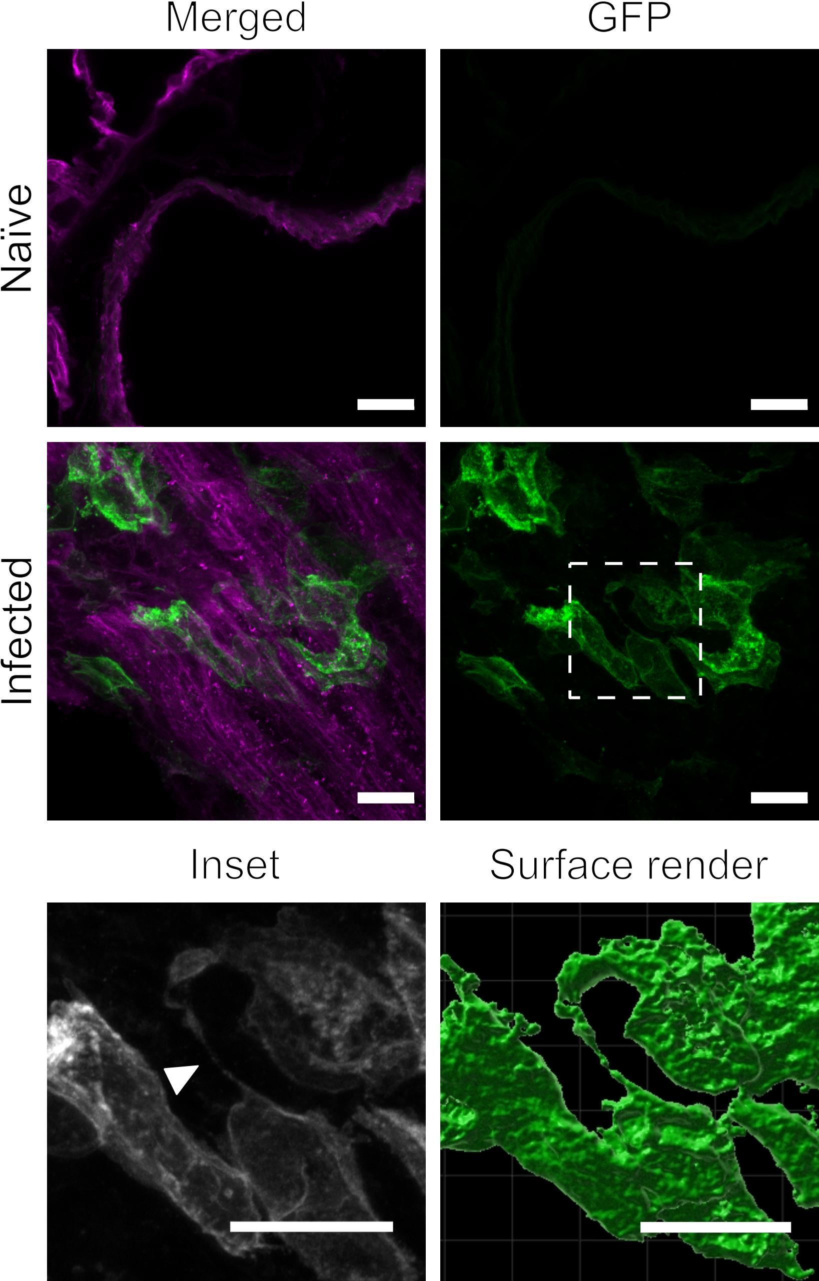

### Supplementary Figure 2

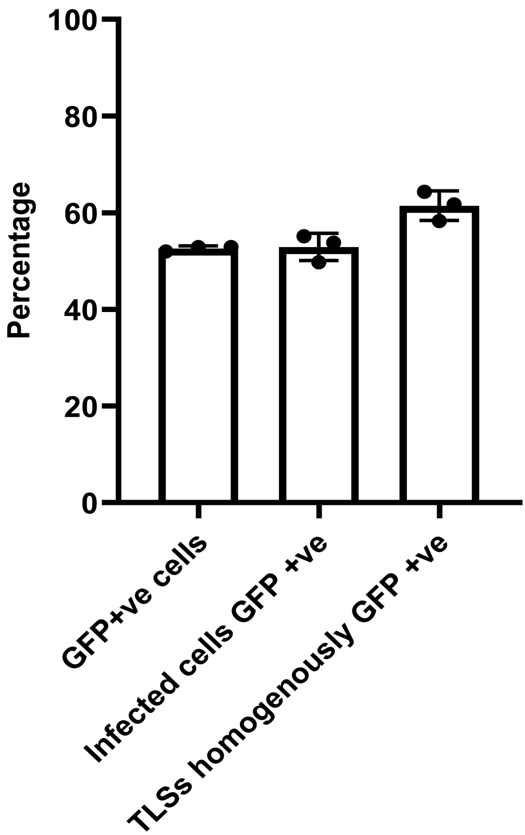

### Supplementary Figure 3

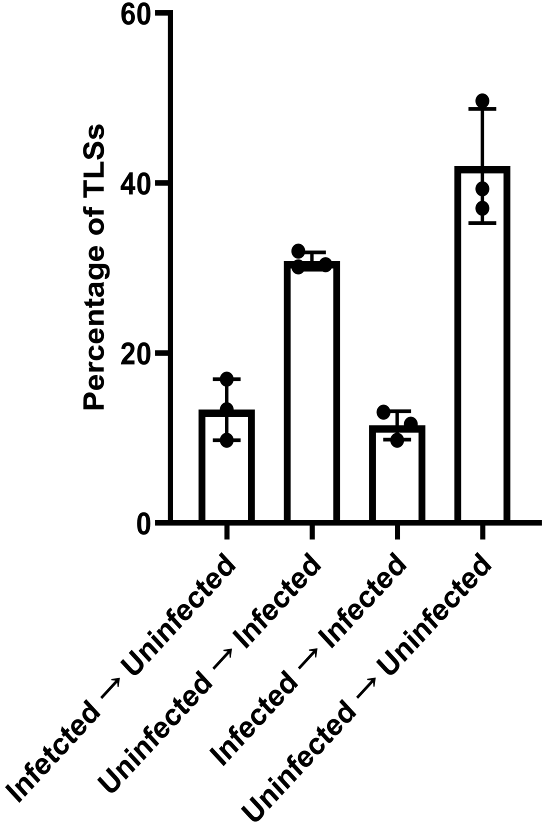

### Supplementary Figure 4

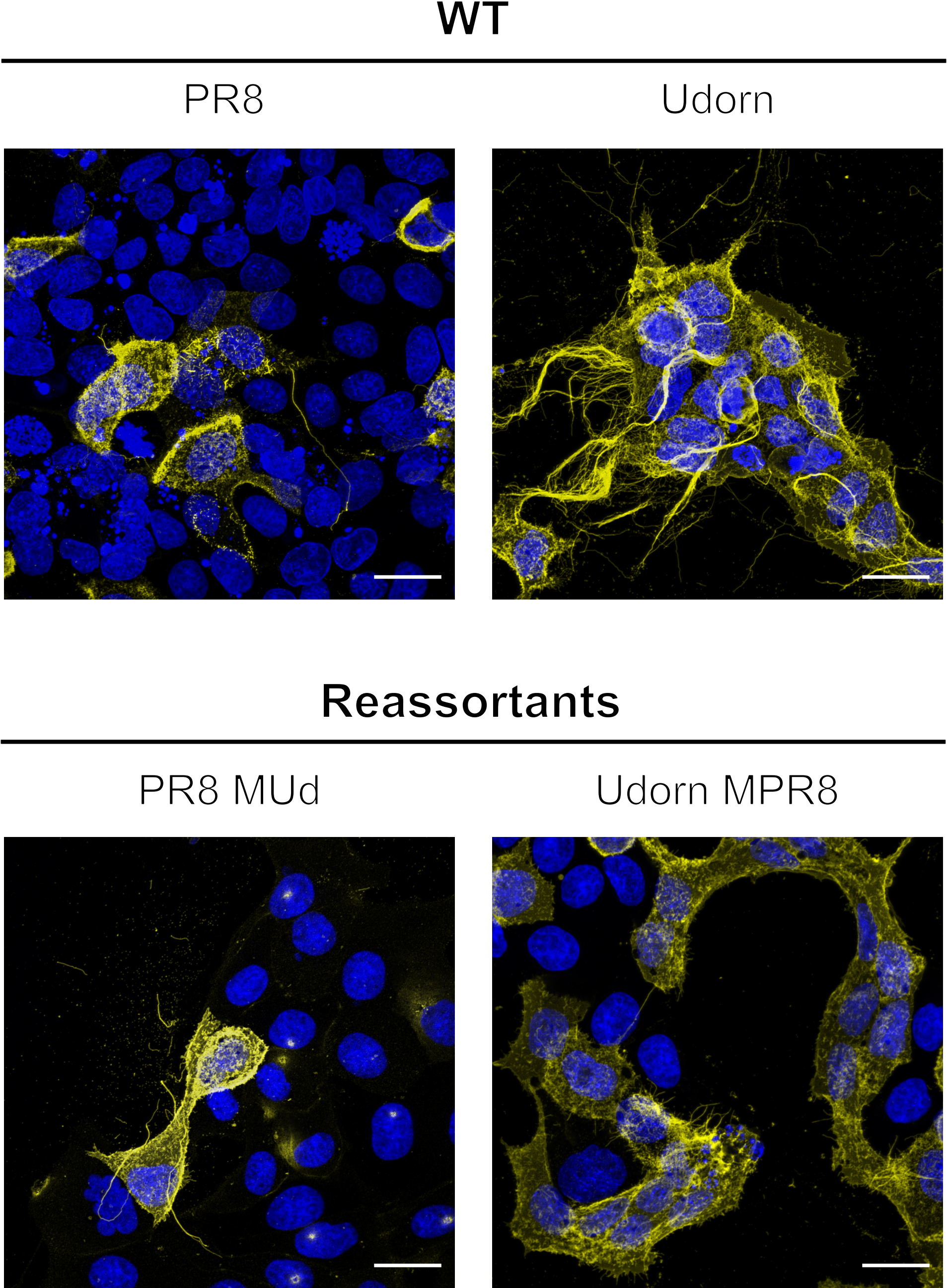

### Supplementary Figure 5

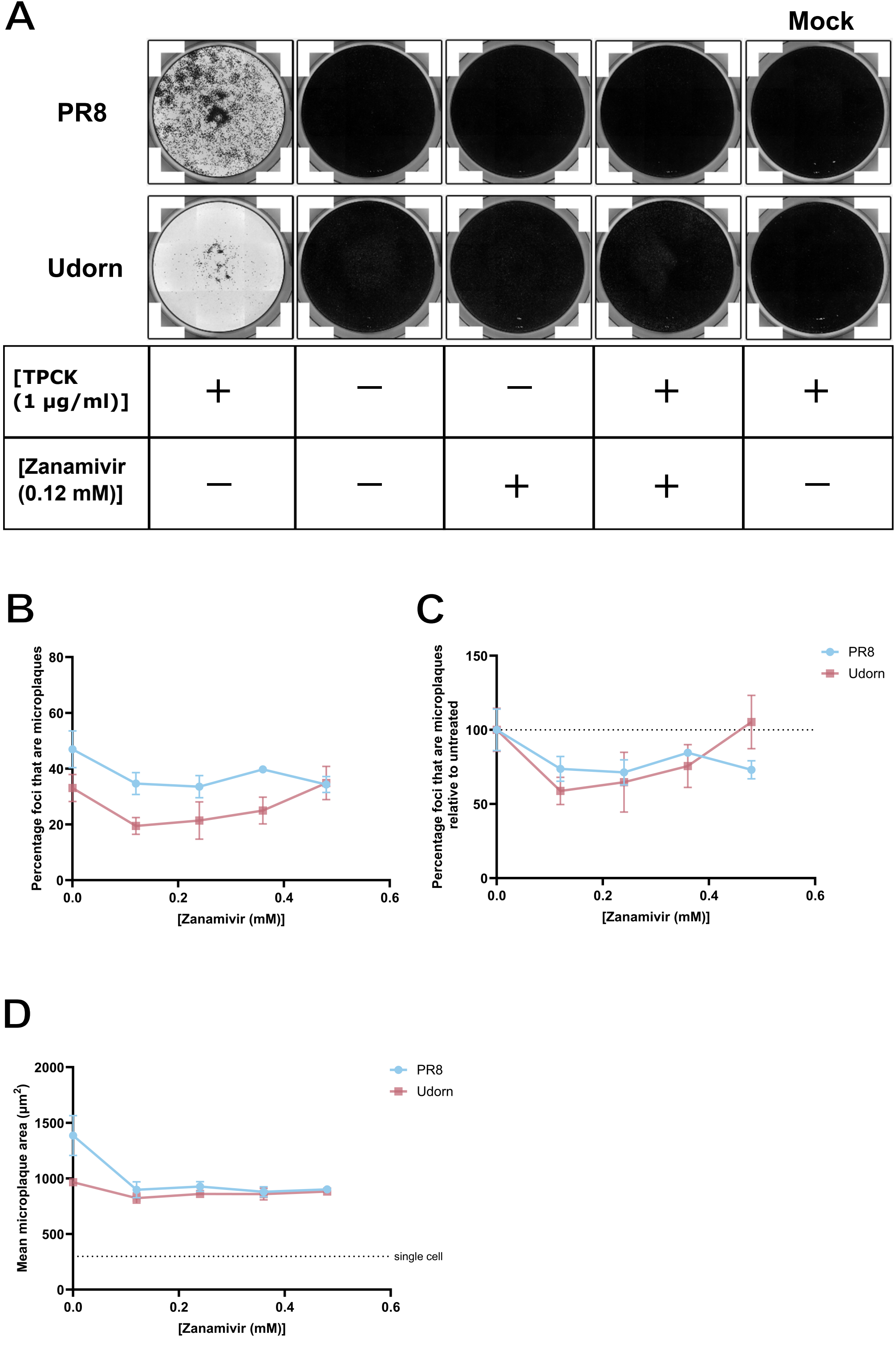
