## Supplementary Figure 6 for "Influenza A viruses induce tunnelling nanotube-like structures through the onset of apoptosis"

# PR8

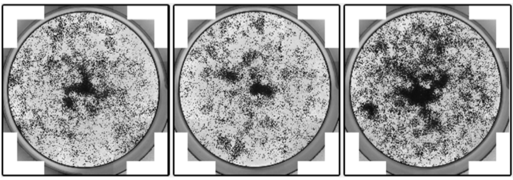

| | | | |
|---|---|---|---|
| [TPCK (1 µg/ml)] | + | + | + |
| Amantadine (mM) | 0.5 | 5 | 50 |

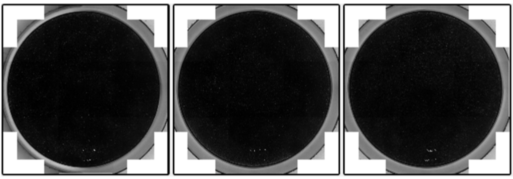

| | | | |
|---|---|---|---|
| [TPCK (1 µg/ml)] | – | – | – |
| Amantadine (mM) | 0.5 | 5 | 50 |

### Udorn

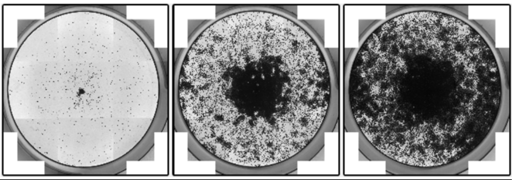

| | | | |
|---|---|---|---|
| [TPCK (1 µg/ml)] | + | + | + |
| Amantadine (mM) | 0.5 | 5 | 50 |

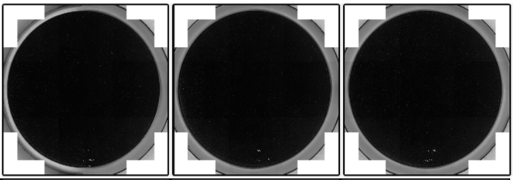

| | | | |
|---|---|---|---|
| [TPCK (1 µg/ml)] | – | – | – |
| Amantadine (mM) | 0.5 | 5 | 50 |
